# Global Spatial Transcriptome of Macaque Brain at Single-Cell Resolution

**DOI:** 10.1101/2022.03.23.485448

**Authors:** Ao Chen, Yidi Sun, Ying Lei, Chao Li, Sha Liao, Zhifeng Liang, Feng Lin, Nini Yuan, Mei Li, Kexin Wang, Meisong Yang, Shuzhen Zhang, Zhenkun Zhuang, Juan Meng, Qiong Song, Yong Zhang, Yuanfang Xu, Luman Cui, Lei Han, Hao Yang, Xing Sun, Tianyi Fei, Bichao Chen, Wenjiao Li, Baoqian Huangfu, Kailong Ma, Zhao Li, Yikun Lin, Zhen Liu, He Wang, Yanqing Zhong, Huifang Zhang, Qian Yu, Yaqian Wang, Zhiyong Zhu, Xing Liu, Jian Peng, Chuanyu Liu, Wei Chen, Yingjie An, Shihui Xia, Yanbing Lu, Mingli Wang, Xinxiang Song, Shuai Liu, Zhifeng Wang, Chun Gong, Xin Huang, Yue Yuan, Yun Zhao, Zhenhui Luo, Xing Tan, Jianfeng Liu, Mingyuan Zheng, Shengkang Li, Yaling Huang, Yan Hong, Zirui Huang, Min Li, Ruiyi Zhang, Mengmeng Jin, Yan Li, Hui Zhang, Suhong Sun, Yinqi Bai, Mengnan Cheng, Guohai Hu, Shiping Liu, Bo Wang, Bin Xiang, Shuting Li, Huanhuan Li, Mengni Chen, Shiwen Wang, Qi Zhang, Weibin Liu, Xin Liu, Qian Zhao, Michael Lisby, Jing Wang, Jiao Fang, Zhiyue Lu, Yun Lin, Qing Xie, Jie He, Huatai Xu, Wei Huang, Wu Wei, Huanming Yang, Yangang Sun, Muming Poo, Jian Wang, Yuxiang Li, Zhiming Shen, Longqi Liu, Zhiyong Liu, Xun Xu, Chengyu Li

## Abstract

Global profile of gene expression at single-cell resolution remains to be determined for primates. Using a recently developed technology (“Stereo-seq”), we have obtained a comprehensive single-cell spatial transcriptome map at the whole-brain level for cynomolgus monkeys, with ∼600 genes per cell for 10 μm-thick coronal sections (up to 15 cm^2^ in size). Large-scale single-nucleus RNA-seq analysis for ∼1 million cells helped to identify cell types corresponding to Stereo-seq gene expression profiles, providing a 3-D cell type atlas of the monkey brain. Quantitative analysis of Stereo-seq data revealed molecular fingerprints that mark distinct neocortical layers and subregions, as well as domains within subcortical structures including hippocampus, thalamus, striatum, cerebellum, hypothalamus and claustrum. Striking whole-brain topography and coordinated patterns were found in the expression of genes encoding receptors and transporters for neurotransmitters and neuromodulators. These results pave the way for cellular and molecular understanding of organizing principles of the primate brain.

## Introduction

Brain is a complex organ comprising a large number of diverse cell types (Hodge et al., 2019; Kebschull et al., 2020; Khrameeva et al., 2020; Krienen et al., 2020; Lei et al., 2020; Network et al., 2020; Tasic et al., 2018; Yuste et al., 2020; Zhu et al., 2018), interconnected to form specific neural circuits that are responsible for animal cognition and behavior (Luo, 2020). Species with higher cognitive and social abilities tend to have larger brains and more neurons relative to the body size (Herculano-Houzel, 2016; Sousa et al., 2017). The macaque brain is composed of more than six billion cells, belonging to hundreds of cell types, each with distinct molecular, morphological, or physiological features (Bakken et al., 2016; Bernard et al., 2012; He et al., 2017; Lein et al., 2017; Maynard et al., 2021). These diverse cell types are located in hundreds of distinct brain regions (Saleem and Logothetis, 2007), which are often defined by anatomical, histological and functional characteristics (Markov et al., 2013). Macaque monkey is a species close to humans and has a greatly expanded neocortex (Singer, 2019), especially the prefrontal cortex (Fuster, 1997; Wise, 2012). Understanding its brain organization at the cellular level holds the key to deciphering neural circuit functions of the human brain and to developing treatments for brain disorders (Buffalo et al., 2019; Eng et al., 2019; Izpisua Belmonte et al., 2015; Lein et al., 2017; Maniatis et al., 2019; Packer and Trapnell, 2018; Stahl et al., 2016; Zhuang, 2021).

Recent advances in single-cell transcriptome analyses have uncovered gene expression profiles of diverse cell types in many brain regions of various species (Bakken et al., 2021; Berg et al., 2021; Khrameeva et al., 2020; Krienen et al., 2020; Lei et al., 2020; Lein et al., 2017; Network, 2021; Yao et al., 2021a; Zhu et al., 2018). Cell types that are unique to mouse and human prefrontal cortex, hippocampus and cerebellum have been found (Aldinger et al., 2021; Berg et al., 2021; Lui et al., 2021; Yao et al., 2021b). The gene expression patterns for various cell types in the developing mouse and human brains have also been characterized (La Manno et al., 2021; Zhu et al., 2018). Comparative studies revealed changes in neuronal types throughout evolution (Bakken et al., 2021; Hodge et al., 2019; Zhu et al., 2018). In addition to the cell type identification in the brain, newly emerged spatial transcriptome technologies have provided new insights into the spatial profiles of gene expression in the brain (Chen et al., 2015; Eng et al., 2019; Lee et al., 2014; Liu et al., 2020; Rodriques et al., 2019; Stahl et al., 2016; Stickels et al., 2021; Vickovic et al., 2019; Wang et al., 2018), revealing cellular organizations that form the basis for the structure and function of neural circuits in the brain. Due to the large brain size and difficulty in retaining cellular organization during tissue preparation, spatial transcriptome analysis over the whole primate brain at single-cell resolution has not been achieved previously. In this study, we employed a newly developed large-scale spatial transcriptome technology Stereo-seq (Chen et al., 2021) and an improved method for obtaining large and thin brain sections. Together with cell type identification by the large-scale snRNA-seq method, we obtained a comprehensive 3-D single-cell spatial transcriptome and cell type atlas for the cynomolgus monkey brain.

The transcriptome datasets here uncovered molecular fingerprints that mark the subregions of the neocortex and subcortical structures, including hippocampus, thalamus, striatum and cerebellum. In addition, we have characterized in more detail spatial single-cell transcriptome profiles in different subdomains of the hypothalamus and claustrum, and mapped the topography of cell type organization in these two structures. Furthermore, we found that the gene expression for neurotransmitter/neuromodulator receptors and transporters exhibited striking topography and coordinated patterns at the whole-brain level, implicating functional linkage of these genes in a region-specific manner. Taken together, our results offer new resources for gene expression profiles at the cellular resolution and whole-brain level, and pave the way for further understanding the organizing principles of the primate brain.

### Spatial transcriptome of entire monkey brain at single-cell resolution

The Stereo-seq (Chen et al., 2021) and droplet-based snRNA-seq DNBelab C4 (Liu et al., 2019) methods were used to obtain single-cell spatial gene expression profile across the monkey brain. A total of 126 coronal sections (10-μm thick, 500-μm spacing) were obtained from the left hemisphere of a freshly isolated and frozen macaque brain (from monkey m1). The 10-μm section thickness allowed the single-cell resolution along the z-axis. We performed brain-wide analysis in 20 coronal sections (CS1-CS20, marked red in Figure 1A-b) at 3-mm spaced coordinates (C1-C20, blue lines in Figure 1A-a), covering the great majority of brain regions (140 out of 143 cortical regions, as defined in (Saleem and Logothetis, 2007); Table S1). For biological replication, we also performed detailed analysis on adjacent sections (CS21-CS25) at five representative coronal coordinates (C3, C6, C11, C13 and C16) from m1, as well as on one 10-μm section each from m2 and m3 (CS26 and CS27 at coordinates C16 and C6, respectively). To obtain high-resolution spatial transcriptome profiles in some neocortical (V1, V2, 44 and 45b) and subcortical regions (hypothalamus and claustrum), we also analyzed a total of 53 brain sections at 500-μm spacing that contained these regions.

**Figure 1.**
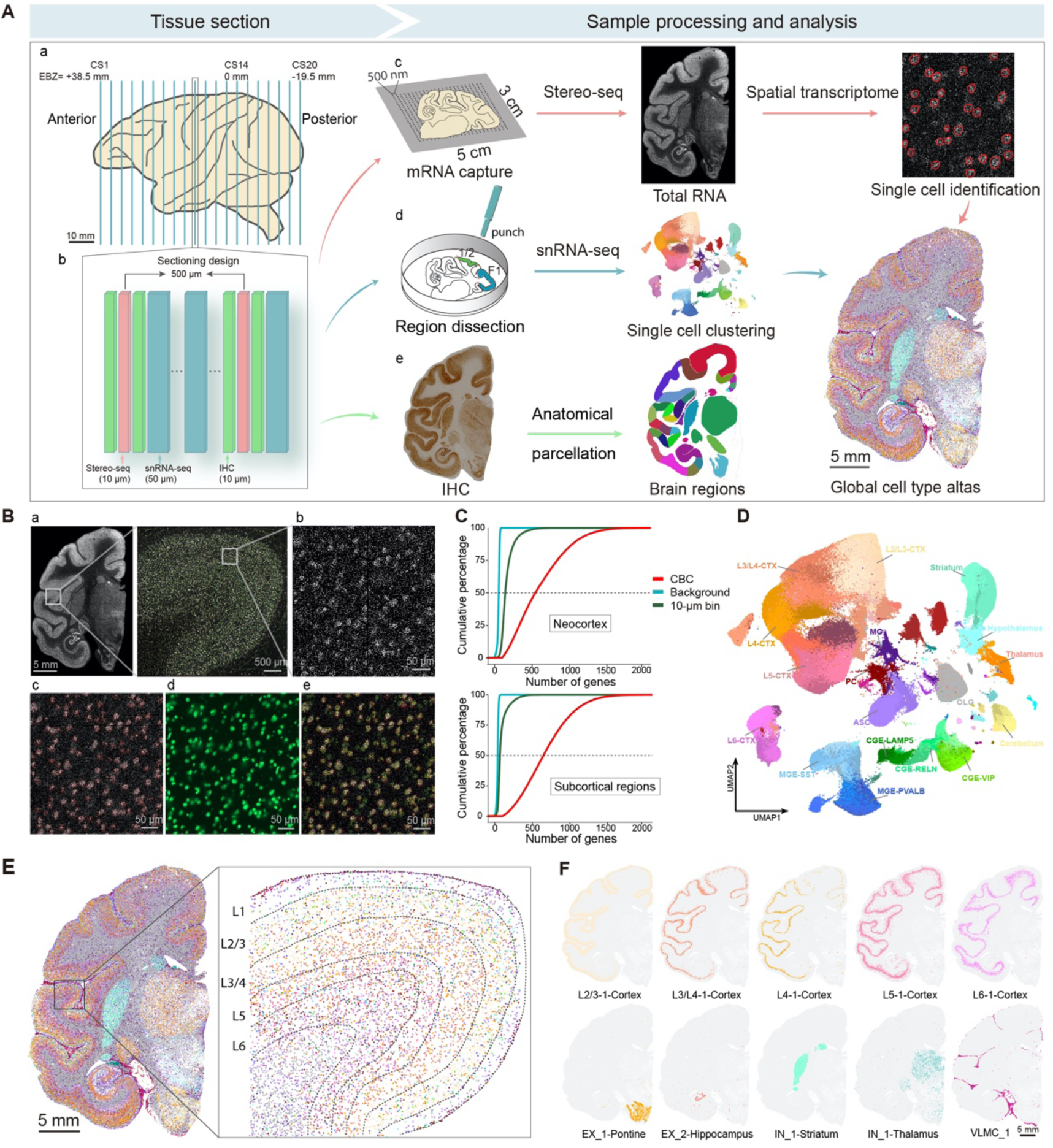
Paradigm for obtaining single-cell spatial transcriptome using Stereo-seq and snRNA-seq. **(A-a)** Schematic diagram showing sample preparation, processing and analysis in this study. **(A-b)** Three types of coronal sections were obtained at each coordinate of the left hemisphere of the monkey brain, consisting of one 10-μm section (red, 3-mm spaced) used for Stereo-seq, two adjacent 10-μm sections for IHC (green) and multiple (3-5) 50-μm thick sections for snRNA-seq analysis (blue). **(A-c)** Stereo-seq sections were used for mRNA capture and spatial transcriptome profiling with single cell identification by CBC method. **(A-d)** Multiple thick sections were micro-dissected for snRNA-seq with region identification and used for cell type clustering analysis and cell annotation to obtain single-cell Stereo-seq transcriptome map. **(A-e)** IHC staining of NeuN for anatomical parcellation of brain regions, marked in color for illustration. **(B)** Illustration of correlation-based circling (CBC) method for single-cell identification in Stereo-seq data. **(B-a)** Total RNA captured from an example section CS10. Area marked with white box is shown in the higher resolution on the right. **(B-b)** The total RNA distribution at cellular resolution in the area marked in red box in **B-a**. **(B-c)** Cells identified by aggregated RNA using CBC method, marked by red dots. **(B-d)** Staining for ssDNA corresponding to the region as in **B-b**. **(B-e)** Merged image of the **B-b, B-c** and **B-d**, showing the high correlation of CBC marked cells and ssDNA staining. **(C)** Cumulative percentage plots of the distribution of the number of genes captured per cell for all cells identified by CBC compared to those of 10-μm binned areas across the entire chip (“10-μm bin”) and outside the CBC-marked areas (“background”) for the neocortex (above) and thalamus (below). **(D)** Cell type identification using clustering analysis of snRNA-seq data on microdissected tissues with region identity. Nineteen supertypes are labeled and 98 different cell clusters are color coded with detailed list in Figure S2A. List of cell type abbreviations is in Table S3. **(E)** An example image for CS10 depicting spatial map of different cell types, color coded as in **D**. Black box on the right depicts cell compositions in the same box area as in **B-a**. Each dot represents a single cell in Stereo-seq data based on the CBC method, with cell identity determined by the similarity between Stereo-seq and snRNA-seq expression profiles. **(F)** Images show the spatial map of ten example cell types in CS10 (color coded as in **D**).

The Stereo-seq method captured mRNAs from all cells of a section in contact with a large silicon chip (up to 5 x 3 cm^2^), which comprises regular arrays of DNA nanoballs (DNB, 220 nm in size, 500 nm center-to-center distance), with each DNB uniquely DNA-barcoded for its spatial location and ligated to polyT sequences for capturing mRNAs from the section. We developed a section flattening method after cryo-sectioning to obtain even and smooth surface contact of large monkey brain sections with Stereo-seq chips (see Methods). To achieve high resolution and spatially resolved cell-type identification, we also obtained multiple 50-μm thick slices (marked blue in Figure 1A-b) adjacent to Stereo-seq sections for conventional snRNA-seq analysis. These slices were dissected with specified region identity for later cell type annotation (Figure 1A-d). For anatomical parcellation of brain regions and verification of gene expression, we also obtained 10-μm sections (marked green in Figure 1A-b) adjacent to Stereo-seq sections for immunohistochemistry (IHC) staining (for NeuN and GFAP; Figures 1A-e and S1A, B).

Due to the high spatial resolution in mRNA capturing capability, the mRNAs captured by Stereo-seq exhibited cell-shaped aggregates (Figures 1B and S1C). We thus developed a <u>c</u>orrelation-<u>b</u>ased <u>c</u>ircling (CBC) method for registering captured mRNAs to corresponding cells (see Methods). The resulting locations of single-cell mRNAs overlapped well with those revealed by the staining of single-strand DNAs (ssDNA; Figures 1B and S1C). The average number of genes obtained in each cell ranged between 100-2000 (see Methods; 663 ± 334 genes/cell), which was 5-fold higher than that obtained by averaging the number of mRNAs collected from all 10-μm bins (20 DNB x 20 DNB, 1-cell size; 146 ± 92 genes/bin, Figure 1C) and much higher than that of the background (Figure 1C). Moreover, CBC-derived single-cell gene expression patterns highly correlated with those from snRNA-seq data of neighboring tissue slices (averaged *R* = 0.90; Figure S1D). Therefore, the Stereo-seq/CBC analysis could reveal spatial gene expression profiles at the single-cell level.

Conventional snRNA-seq analysis was performed on dissected tissues from multiple (15-30) 50-μm thick slices adjacent to the corresponding Stereo-seq sections (see Methods). Transcriptional profiles of a total of 652,225 cells (2,990 ± 1,132 genes per cell) were obtained after quality control (Table S2). Dimensionality reduction followed by clustering analysis of snRNA-seq data (see Methods) led to the identification of 98 cell clusters (Figures 1D and S2A). Based on previously known cell-type marker genes (Ximerakis et al., 2019; Yuste et al., 2020; Zeisel et al., 2018) and region identity information obtained during dissection, we assigned the 98 clusters into 4 classes and 18 supertypes (Figure 1D). The excitatory neuron class contains 5 supertypes (L2/3-CTX, L3/4-CTX, L4-CTX, L5-CTX and L6-CTX) and 27 clusters, and inhibitory neuron class contains 5 supertypes (CGE-LAMP5, CGE-VIP, CGE-RELN, MGE-SST, and MGE-PVALB) and 29 clusters. Each of the non-neuronal and subcortical classes contains 4 supertypes (Figure S2A). These cell types expressed known marker genes, e.g., *SLC17A6*, *SLC17A7*, *GPR83*, *CUX2*, *RORB*, *ETV1* and *SEMA3E* for excitatory neurons, *GAD1*, *GAD2*, *RELN*, *SST, PVALB, VIP and LAMP5* for inhibitory neurons, *GFAP* and *S100B* for astrocytes, *ITGAM* for microglia cells, and *MOBP* for oligodendrocytes (Figure S2B).

We annotated the single cells obtained by Stereo-seq, based on the highest correlation of their gene expression profiles with those of 98 different cell types determined by snRNA-seq (see Methods; Figure S3A). We then generated comprehensive spatial cell maps, comprising over 3 million single cells (60,227 – 310,211 cells per section), across 20 coronal sections (Figures 1E and S3B). Such single-cell spatial map revealed many fine details of brain structures including neocortical layers, white matter, and distinct domains of subcortical areas. Figure 1E shows a composite image of the distribution of 98 cell types in CS11, together with a zoomed-in image showing their laminar distribution of cell types. In addition, specific excitatory and inhibitory neuron types were found in subcortical areas, and non-neuron cell type VLMC was mainly distributed in the pial surface (Figure 1F). Similar cell type-specific spatial maps were also observed in the adjacent sections from the same monkey and corresponding sections from two other monkeys (Figure S3B). These results indicated the feasibility and reliability of our spatial map of single cells by combining Stereo-seq and snRNA-seq methods.

To proceed towards the construction of the global 3D cell type atlas for the entire monkey brain, we performed combined analysis of Stereo-seq and snRNA-seq data for all 20 coronal sections (3-mm spacing), as shown in Figure 2A. The overall cell-type distributions exhibited consistent patterns and gradual changes among adjacent sections, and transcriptome patterns between neocortical vs. subcortical regions were drastically different in all sections (Figure 2A). Figure 2B illustrates the side view of the global distribution for four layer-specific neuron subtypes in the neocortex that exhibited striking specificity along the anterior-posterior axis of the elongated brain (3D visualization in Video S1). Figure 2C illustrates the side view of the global distribution for several types of excitatory neurons, inhibitory neurons, and astrocytes that were found only in specific subcortical structures such as hippocampus, thalamus, striatum or cerebellum (3D visualization in Video S2). Cell type compositions in various neocortical and subcortical structures were quantified over the entire monkey brain. The percentage of various cell types in each region was quantified and plotted in the heatmap (Figure 2D, more detailed information in Figure S3C and Table S3). The 3D cell type atlas obtained here, together with quantification of their global distribution and transcriptomic profiles, provides a useful resource for further analysis of the cellular organization of the primate brain.

**Figure 2.**
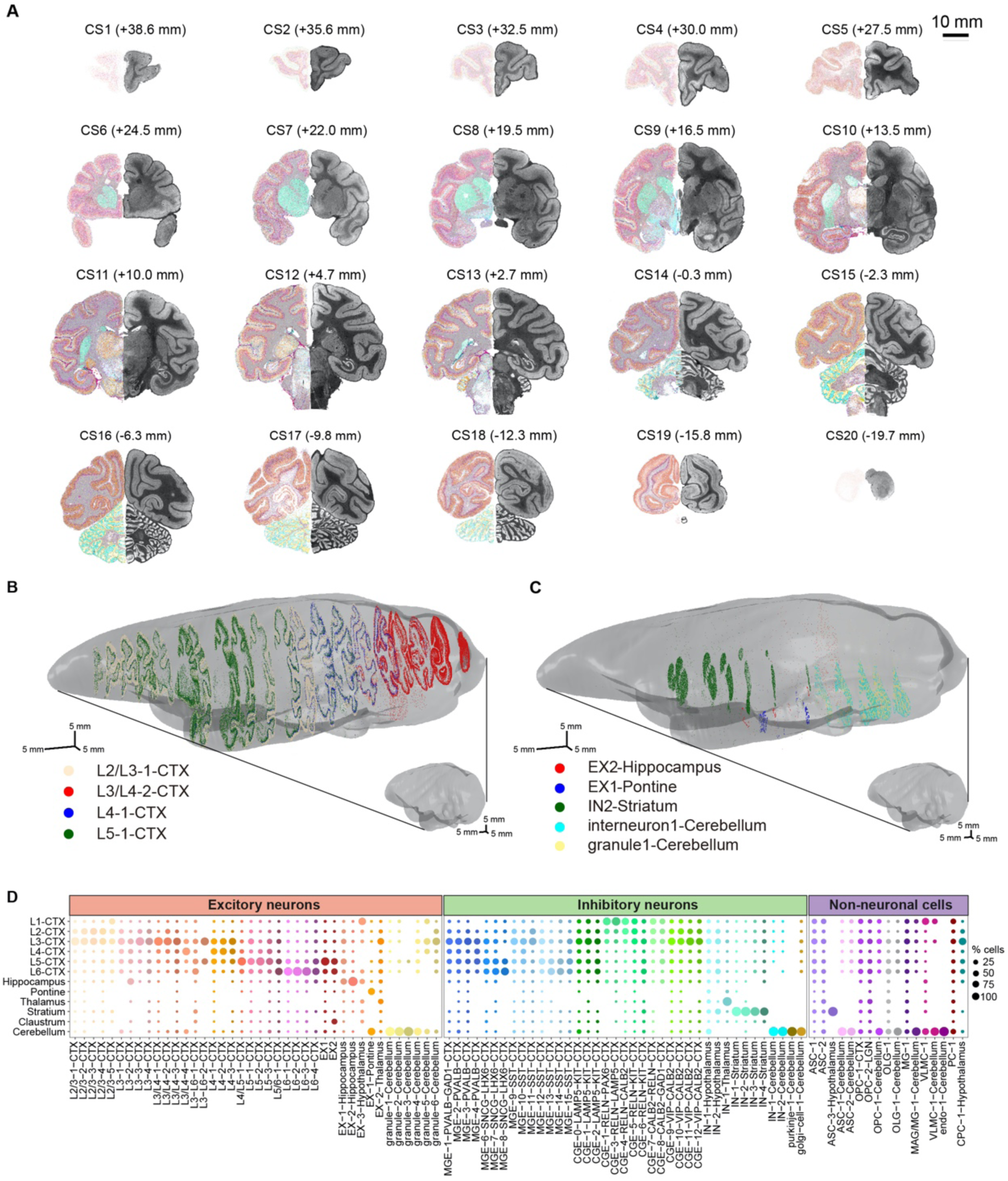
Global transcriptome map of monkey brain at single-cell resolution. **(A)** Spatial map of different cell types for 18 coronal sections across the entire monkey brain, with cell types color coded as in Figure 1D and AP coordinates shown in parenthesis. Composite cell-type distribution and raw total mRNA map of the same left-hemisphere section were presented together as mirror images. **(B-C)** Side view of the 3D map of selected neocortical **(B)** and subcortical **(C)** cell types in the monkey brain elongated along the AP axis. **(D)** Cell type composition of different neocortical layers and subcortical structures at the whole brain level revealed by Stereo-seq analysis (color coded as in Figure 1D). Dot size represents the percentage distribution of each cell type in various cortical layers and subcortical structures, with the sum of each column equals to 100%.

### Neocortical layer- and region-specific gene expression profiles

A hallmark of the neocortex is the laminar organization of neurons with distinct connectivity and functional roles. We thus further examined the laminar specificity in cell-type distribution, which could be readily identified in the Stereo-seq maps (Figures 1E, 1F and 2A). Most excitatory neuron types showed some layer preferences (Figures 1E, F and 2D), consistent with previous findings (Ximerakis et al., 2019; Zeisel et al., 2018). Inhibitory neurons in general showed lower layer preferences, except for *reelin*-expressing CGE-RELN neurons (Rice and Curran, 2001), which were predominantly located in L1 and L2/3 (Figure 2D).

We further characterized layer-specific genes identified by Stereo-seq across the entire monkey brain, by analyzing differential gene expression across NeuN immunostaining-defined cortical layers (see Methods). Examples of layer-specific expression of the top1-ranked genes for each layer are shown for five typical coronal sections, with each color depicting the expression level of one example gene (CS11 in Figure 3A and others in Figure S4A). In addition to the genes previously reported(Bakken et al., 2016), our Stereo-seq data revealed many new genes showing layer-specific expression in multiple cortical areas (Figures 3B and S4B, C). For example, *FABP7* in L1, *GPR83* in L2, *ADCYAP1* in L3, *IL1RAPL2* in L4, *ETV1* in L5, and *KRT17* in L6. Some layer-specific genes showed widespread distribution in many cortical regions (Figures 3B and S4B), while others showed more restricted regional specificity (e.g., *NEFH* in L5 of F1 and *OSTN* in L4 of V1; Figure S4C). We have identified 219 genes on average in each cortical area (2,638 genes in total) showing layer-specific expression (see Methods; Table S4). These layer-specific genes were validated in adjacent sections (500 μm away, CS21-25) from the same monkey (m1) and corresponding sections (CS26-27) from two other monkeys (m2 and m3; Figure S4).

**Figure 3.**
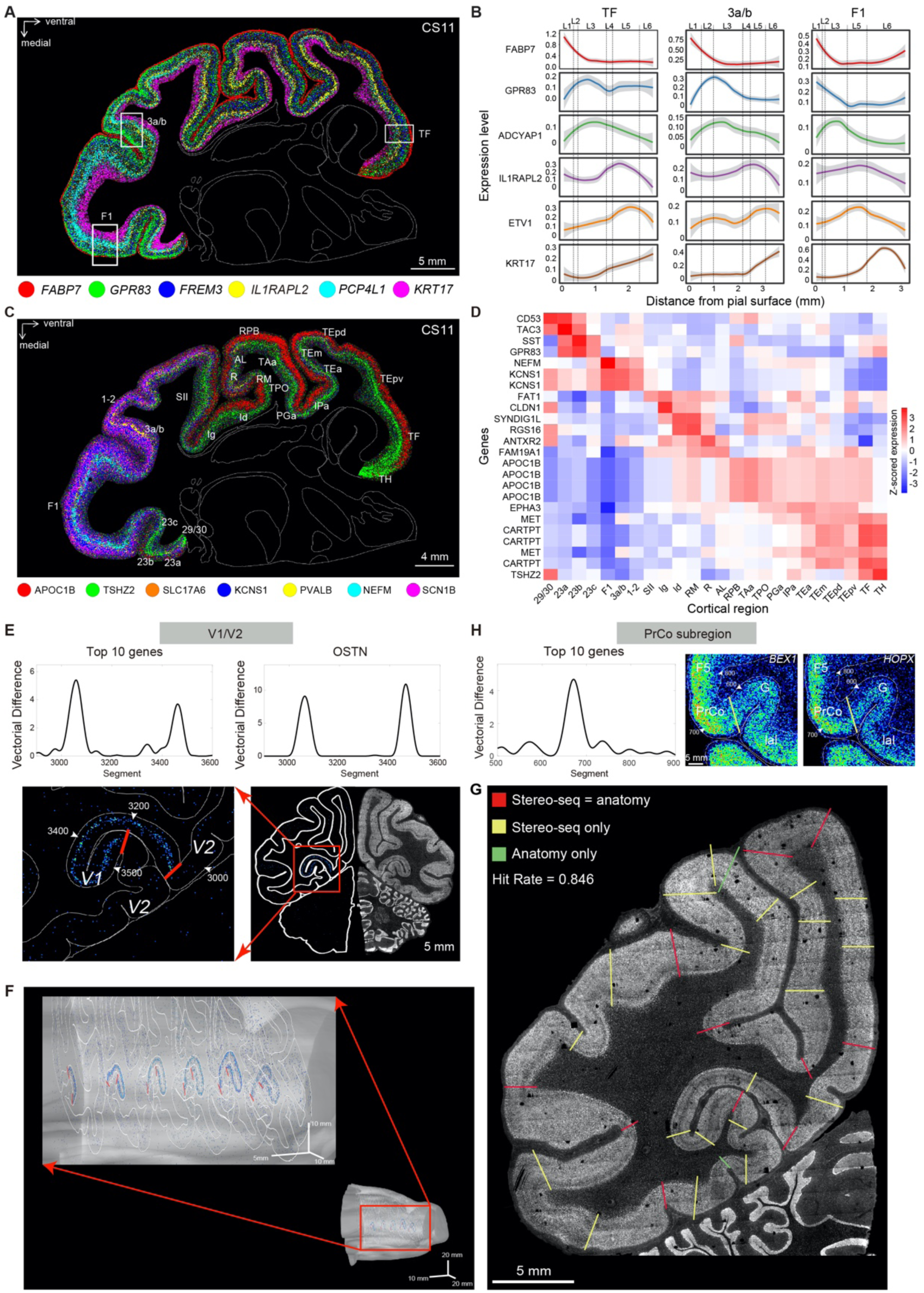
Layer- and region-specific gene expression profiles and their utility in brain parcellation. **(A)** Composite image of expression profiles for 6 example layer-specific genes across the coronal section CS11, with each gene coded with a distinct color and its expression level (summed in 25-μm bins) coded by color intensity. **(B)** The expression profiles for the 6 example layer-specific genes in **A** were quantified for 3 different brain regions marked by arrowheads in **A**. **(C)** Composite image of expression profiles for 6 example region-specific genes in CS11. Region-specific genes are coded by colors the same manner as in **A**. **(D)** Heatmap showing the expression level of 25 genes, each was ranked top in the expression level for a specific region in section CS11. The expression level was z-score normalized for each gene across the 25 cortical regions. **(E)** Regional boundary identification by gene expression profiles, as illustrated for the boundary between V1 and V2. Top curves represent vectorial differences of gene expression between adjacent cortical segments (see text), illustrated for averaged top 10 genes (top left) and an example gene *OSTN* (top right) for the V1/V2 boundaries. Images below showed spatial expression patterns of the *OSTN* gene along the cortical sheet, red lines indicate the V1/V2 boundaries, arrowheads mark the segment numbers corresponding to the profile above. **(F)** Anterior-posterior elongated 3D map showing V1/V2 boundaries across 6 coronal sections (500-μm spacing) of the m1 monkey brain. **(G)** A new boundary identified in PrCO region using same analysis as in **E**. **(H)** Image of total mRNAs for coronal section CS25, showing regional boundaries that were marked by gene expression patterns alone (yellow), anatomy alone (green), and by both (red).

We then quantified the total expression level of each gene across the neocortex and found 3,650 genes that exhibited region-specific expression (see Methods; Figure S5A and Table S5). Figure 3C shows a composite image of expression patterns of seven region-specific genes in CS11. Some genes showed distinctly higher expression in one or a few regions (*NEFM* in F1, *KCNS1* in 1-2 and 3a/b), while others (*APOC1B*, *TSHZ2*, *SCN1B, PVALB* and *SLC17A6*) were expressed more widely in many regions. An average of 83 genes with region-specific expression were found in each cortical area (Table S5), and top-ranked genes were listed for various cortical regions (CS11 in Figure 3D, and CS3, CS6, CS13 and CS16 in Figure S5B-E). Besides a few previously reported genes(Ataman et al., 2016) (e.g., *OSTN* in V1 and *NEFM* in F1), we found many genes showing region-specific expression, such as *CENPF, SYT6, CHRNA1, UCMA* (in V1)*, KCNS1* (in F1, 1-2 and 3a/b), *PTGDS* (in V4v), *GDF10, DIAPH3, PCDH17* (in 9m, 9d and 10mr), and *OGN*, *CRABP1* (in TH; Figures 3D, S5B-E and S6). Such region-specific expression is in line with the region-specific composition of cell types (Figure S3C), and was further validated in adjacent sections (500 μm away) of the same monkey (m1) and corresponding sections in the two other monkeys (Figure S6).

Mapping cortical subregions is of major importance to systems neuroscience, but precise parcellation of the cortex has been challenging (Van Essen and Glasser, 2018). We reasoned that information on region-specific gene expression may facilitate cortical parcellation. To search for gene expression patterns that could define subregion boundaries, we quantified the differences in gene expression profiles along the cortical sheet. This was achieved by dividing the cortical sheets into vertical sliding segments (150-μm width, 15-μm step size), and each segment was further divided horizontally and evenly into 20 laminar sub-divisions (see Figure S7A and Methods). The expression level of each gene within a segment was represented by a 20-dimension vector, with the component representing the expression level in each sub-division. The adjacent segmental differences in gene expression were quantified by vectorial differences for all genes (see Methods; Figure S7A). Expression difference curves along the cortical sheet of each section showed peaks representing sharp transitions in gene expression profiles, and the location of these peaks were used to indicate region/subregion boundaries in the cortex (Figure 3E-H). One clear example was the boundaries between V1 and V2 regions (Figure 3E), which were further visualized in 3D plot with all serial sections containing V1/V2 regions (Figure 3F). The genes that provided dominant contributions for defining the V1/V2 boundary were *OSTN, PHACTR2, KCNH8* and *ADAMTS17* (Figure S7B). Another example of clearly defined boundaries was shown between regions 44 and 45b (Figure S7C, D), based largely on the expression profiles of *SCN1B*, *KCNS1*, *TRIM55* and *RING1* (Figure S7C). These boundaries were also validated in adjacent sections (Figure S7B-D).

We found that a majority of anatomically defined regional boundaries coincided with Stereo-seq defined boundaries (84.6% in CS25; Figure 3G, red lines). This was significantly higher than the chance level determined by random shuffling of boundaries (*P* = 0.002; see Methods). Furthermore, some peaks in expression difference curves revealed previously undefined boundaries (Figure 3G, yellow lines), and on average 1.5 new subregion boundaries were found within each anatomically defined region. For example, a new subregion boundary was defined within PrCO (precentral opercular area), based on the expression profiles of *BEX1*, *HOPX*, *RPB4* and *MPC2* (Figure 3H). We noted that substantial fraction (∼15.4% in CS25) of anatomically defined boundaries were not marked by distinct changes in gene expression profiles (Figure 3G, green lines). Thus, spatial transcriptome profiles obtained by Stereo-seq offer a new type of information for parcellating the brain.

### Domain-specific gene expression profiles of subcortical structures

We next analyzed region-specific gene expression profiles in different subcortical regions. Many genes were expressed in a specific manner in various subcortical structures (quantified as in Methods). Table S6 summarizes all region- and domain-specific genes and their expression levels in various subcortical structures examined in this study. For examples, *SYNPR*, *TRPM3*, *PENK*, *TCF7L2*, *SLC18A2* and *CABP7* were highly expressed in the claustrum, choroid plexus, striatum, thalamus, substantia nigra, and pontine nucleus, respectively (Figure 4A and Figure S8A). Previous single-cell transcriptome studies in monkeys and humans have identified a variety of cell types in the hippocampus, thalamus, striatum and cerebellum (Kebschull et al., 2020; Zhong et al., 2020; Zhu et al., 2018), but the spatial distribution of these cell types and gene expression profiles remain to be elucidated. We thus further examined the spatial profiles of gene expression in these structures.

**Figure 4.**
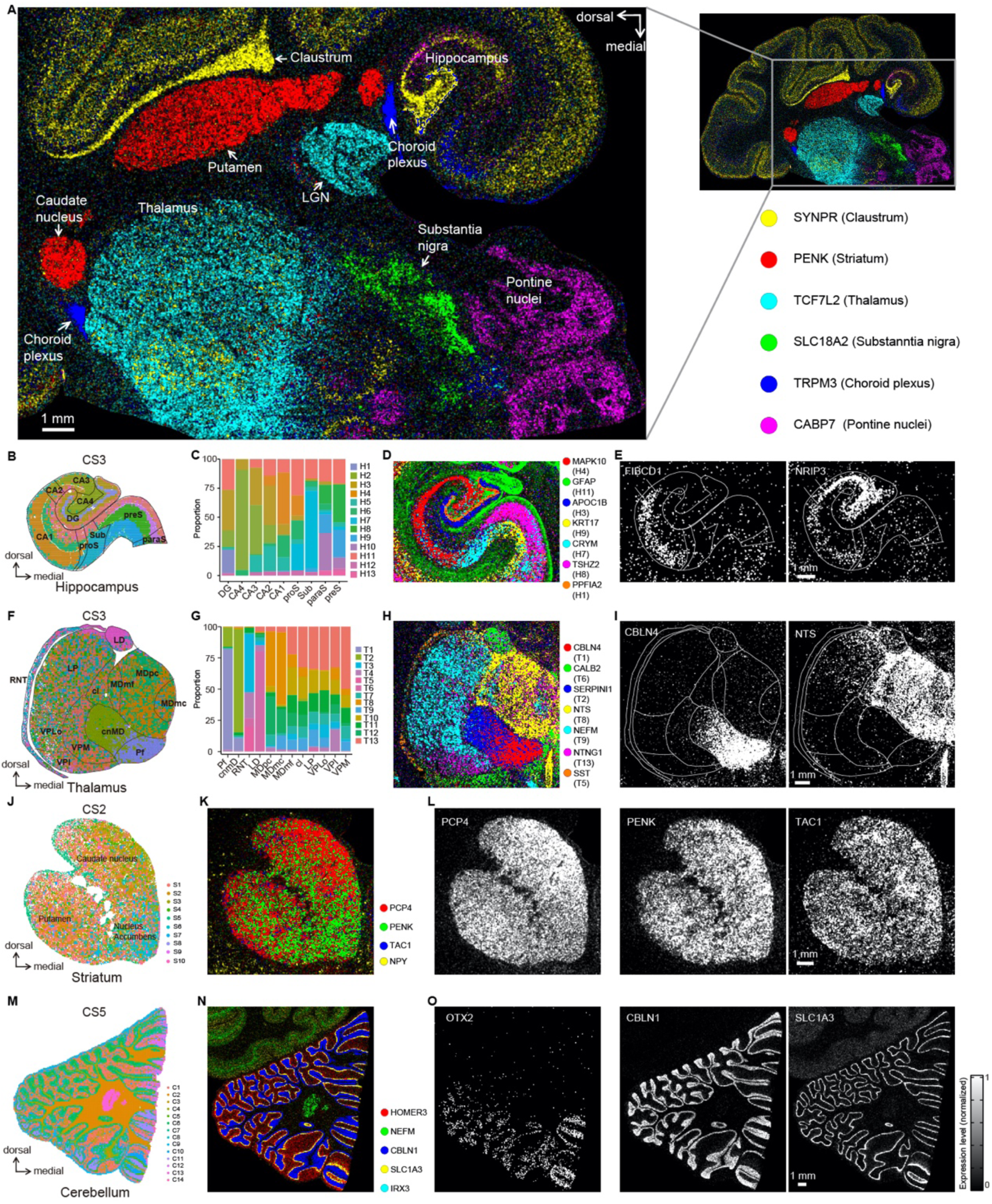
Domain-specific gene expression profiles of subcortical structures. **(A)** A composite image of the expression profiles for domain-specific genes in the subcortical structures of coronal section CS11. Inset: location of subcortical structures relative to the neocortex. Examples of structure-specific genes are color coded. **(B-E)**, Expression profiles of subdomain-specific genes in the hippocampus. Subdomains are marked by black lines in **B**. Colors mark different clusters (1-13) of gene expression profiles as determined by clustering analysis of Stereo-seq data (at 100-μm bins). The percentage of various clusters found in hippocampal subdomains are shown in **C**. The expression profiles for 7 subdomain-specific genes in the hippocampus are shown in the composite image in **D**, and that for 2 example genes are shown separately in **E**. (**F-I)**, Subdomain-specific gene expression profiles in the thalamus, presented similarly as in **B-E**. **(J-L)**, Spatial gene expression profiles in the striatum. Ten clusters of gene expression profiles determined by Stereo-seq data are marked by different colors in **J**. The expression profiles for 4 example genes are shown in the composite image **K** and 3 of them are shown in separate images **L**. **(M-O)**, Spatial gene expression profiles for the cerebellum, with distinct distributions of 14 cell clusters marked in colors in **M**. A composite image of expression profiles for 5 genes and separate images for 3 genes are shown in **N** and **O**, respectively.

By unsupervised clustering analysis of gene expression profiles from Stereo-seq data at 100-μm bins, we identified 13 expression clusters in the hippocampus (see Methods; Figure S8B). Spatial coordinate mapping revealed distinct patterns of spatial localization for each expression cluster (Figure 4B). Notably, these clusters showed clear correspondence to hippocampal domains that were defined by anatomical landmarks and NeuN staining (Figure 4B, C). For examples, more than 70% of cells in clusters H1 and H7 were localized to Dentate Gyrus (DG) and Subiculum (Sub), exhibiting high expression of *PPFIA2* and *CRYM*, respectively (Figure 4B, C and Figure S8C). For different cell types, we also found specific expression of genes that marked different subdomains, as shown in the composite image for seven example genes: the expression of *APOC1B*, *MAPK10*, *CRYM*, *TSHZ2* and *GFAP* was largely confined to DG, Cornu Ammonis (CA) 1-4, Sub, pre-Subiculum (pre-S), and peri-hippocampal area, respectively (Figure 4D). Further examples of two domain-specific genes *FIBCD1* and *NRIP3,* which were largely confined to CA1 and CA2-4, respectively, are shown in Figure 4E. Notably, *NRIP3* localization in CA2-4 was consistent with that found in human hippocampus (Zhong et al., 2020).

Similar clustering analysis based on spatial gene expression profiles in the thalamus, cerebellum and striatum also identified distinct gene expression clusters (Figures 4F-O; top two marker genes shown in Figure S8B, C). Notably, we found four clusters (T1-4) exhibiting clear preference in distinct thalamic subdomains (Figure 4F, G). For examples, the Paratenial nucleus (Pt) and Laterodorsal Nucleus (LD) were mainly (> 80%) composed of cells of cluster T1 and T6, showing elevated expression of *CBLN4* and *CALB2*, respectively (Figure 4F-I). In contrast to the thalamus, gene expression showed less distinct domain specificity in the striatum (Figure 4J-L). For examples, *PCP4* showed higher expression in the dorsal caudate nucleus, and *PENK* and *TAC1* showed patchy expression patterns across the entire striatum (Figure 4L). In the cerebellum, the spatial expression patterns of many genes were markedly different from other brain structures. For examples, we observed a ventral-dorsal decreasing gradient of *OTX2* expression, and striking complementary pattern of *CBLN1* and *SLC1A3* (Figure 4O). These domain-specific gene expression profiles in different subcortical structures described above were validated in adjacent coronal sections (500 μm away) in the same monkey m1 and the corresponding section from monkey m2 (Figure S8A). Distinct genes expressed in specific domains of subcortical structures identified here represent interesting targets for future functional studies.

### Single-cell spatial transcriptome map of monkey hypothalamus

The hypothalamus comprises multiple nuclei and regulates many innate behaviors and homeostatic functions (Andermann and Lowell, 2017; Anderson, 2016; Li and Dulac, 2018; Morrison and Nakamura, 2019; Sternson, 2013; Weber and Dan, 2016; Zilkha et al., 2017). The functional roles of various hypothalamic cell types have been extensively characterized in mice (Andermann and Lowell, 2017; Anderson, 2016; Kim et al., 2019; Li and Dulac, 2018; Mickelsen et al., 2019; Moffitt et al., 2018; Morrison and Nakamura, 2019; Romanov et al., 2017; Sternson, 2013; Weber and Dan, 2016; Zilkha et al., 2017). However, whether these cell types are evolutionarily conserved in monkeys and how they are spatially organized remain largely unknown. Using Stereo-seq data for 12 sections at ∼500 μm spacing (Figure 5A), we analyzed the spatial transcriptome profiles in the monkey hypothalamus at the single-cell resolution. To assist cell type annotation of Stereo-seq data, we performed unsupervised clustering analysis on snRNA-seq data of ∼10,000 cells that were obtained from dissected hypothalamus tissue (Figure S9A). This led to the identification of five major cell classes, including inhibitory and excitatory neurons, astrocytes (ASC), ependymal cells (EPC) and oligodendrocyte progenitor cells (OPC; Figure S9A, B). Further clustering analysis revealed 28 and 13 subtypes within inhibitory neurons (6,400 cells) and excitatory neurons (1,172 cells), respectively (Figure S9C, D). Notably, each excitatory and inhibitory neuron subtype exhibited distinct neuropeptide expression patterns, many were specifically marked by a single neuropeptide (e.g., AVP for EX4, OXT for EX9, HCRT for EX11, AGRP for IN13 and GHRH for IN25), and most exhibited unique combinations of neuropeptides (Figures 5B, C and S9F, G). This supports the notion that unique cell types could serve specific hypothalamic functions through the action of distinct combination of neuropeptides.

**Figure 5.**
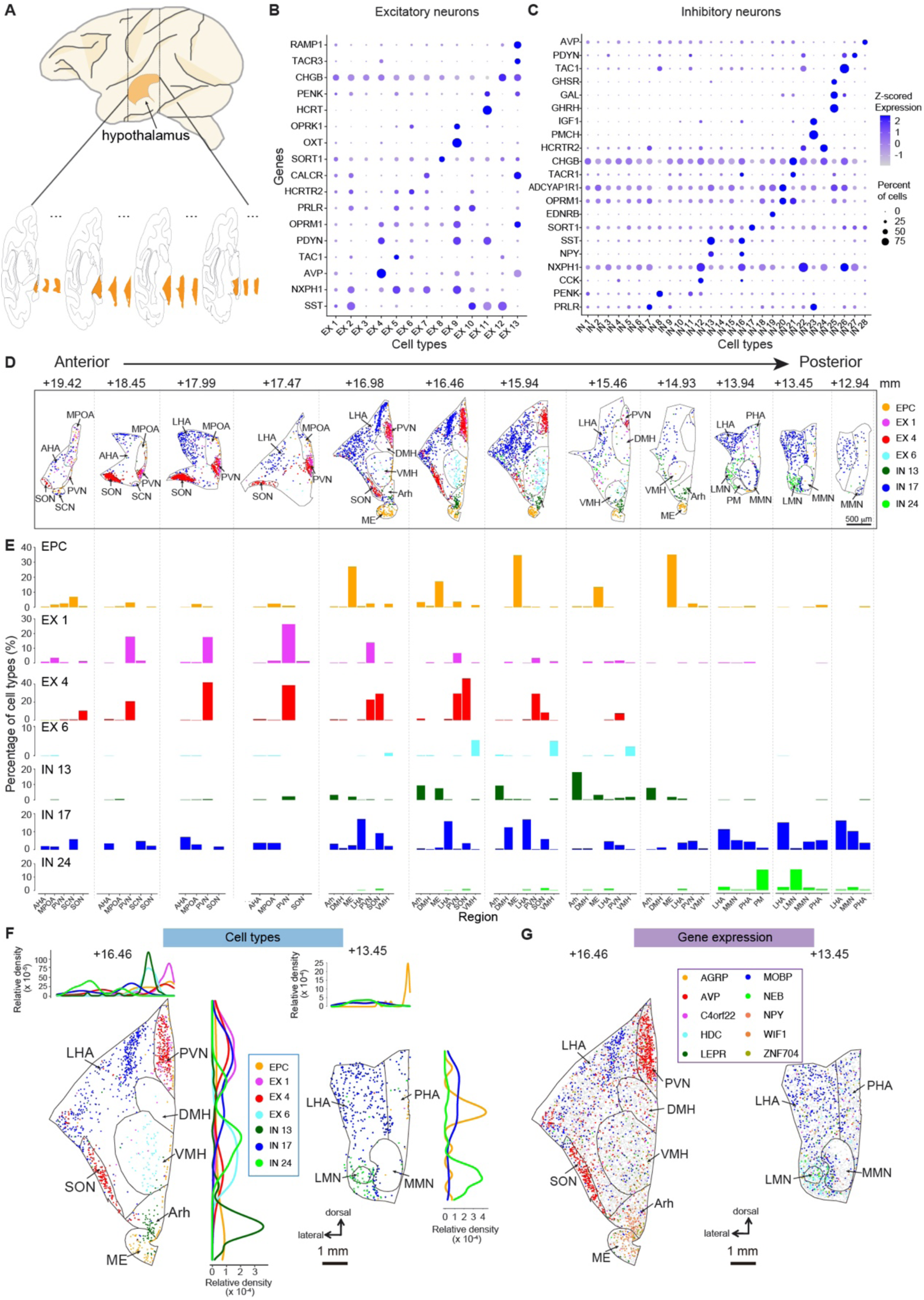
Single-cell spatial transcriptome in monkey hypothalamus. **(A)** Schematic diagram indicating the location of hypothalamus sections of the macaque brain. **(B-C)** Dot plot showing the normalized expression for neuropeptides within excitatory (B) and inhibitory (C) neuron subtypes identified by snRNA-seq data of hypothalamus tissues. The dot size represents the percentage of cells in each subtype expressing the given neuropeptide. The list of neuropeptide abbreviations in Table S7. **(D)** Spatial distribution of 7 example cell types for 12 sections along the anterior-posterior axis of the hypothalamus. Coronal coordinates shown by the numbers in mm above. **(E)** Quantified cell type distribution of 7 example cell types in the identified nuclei of each section shown in D. **(F-G)** Images show the spatial map of selective cell types (F) and marker gene expression (G) at single-cell resolution in an example coronal section AP (+16.46 mm). Density plots above and beside the section image showed spatial distributions of the 7 cell types along the medial-lateral and dorsal-ventral axis, respectively. **(H-I)** As in F and G for another example coronal section AP (+14.93 mm).

We then annotated 47,258 single hypothalamic cells in the Stereo-seq map (Figure S9E), based on the highest correlation of their gene expression profiles with the cell types determined by snRNA-seq analysis above. This provided the spatial distribution of various cell types within the hypothalamus. As shown in Figure S9H, most cell types exhibited distinct distribution in various hypothalamic nuclei that were defined by histological parcellation with NeuN staining. In particular, AVP-expressing EX4 subtype was enriched in supraoptic nucleus (SON) and periventricular nucleus (PVN), whereas NXPH1-expressing EX5 and HCRTR2-expressing EX6 subtypes were enriched in ventromedial hypothalamic nucleus (VMH). On the other hand, inhibitory neuron subtypes, which were more abundant than excitatory neuron subtypes, appeared to distribute in a more dispersed manner. However, some subtypes were enriched in specific nuclei, such as SST/NPY-expressing IN13 and GHRH/GAL-expressing IN25 in arcuate hypothalamic nucleus (Arh), SORT1-expressing IN17 in lateral hypothalamic area (LHA) and HCRTR2-expressing IN24 in lateral mammillary nucleus (LMN; Figures 5D and S9H). For non-neuronal cells, EPCs were enriched in median eminence (ME), while ASCs and OPCs showed wider distribution (Figures 5D and S9H). Comparison of the transcriptome data between mouse and monkey hypothalamus showed both conserved and divergent patterns of gene expression. For example, previous studies in mice showed high expression of AVP and OXT neuropeptides in the excitatory neurons of both SON and PVN (Moffitt et al., 2018; Romanov et al., 2017; Zhang et al., 2021), a pattern that observed here in monkeys. In contrast, AVP-expressing inhibitory neurons were enriched only in mouse SCN (Moffitt et al., 2018), but were also present in monkey SON and ME.

Further examination revealed clear preferences of several cell types along the anterior-posterior axis of the hypothalamus, consistent with the localization of individual cell types to different nuclei. Notably, EX1 and EX4 neuron subtypes were enriched in the anterior hypothalamus, EX6, EPC and IN13 subtypes were distributed mainly in the medial region, whereas IN17 and IN24 subtypes were enriched in the medial-posterior region (Figure 5D, E). In addition, we found spatial preferences of different cell types along the dorsal-ventral and medial-lateral axis of the hypothalamus (Figure 5F). For examples, EPC and IN13 types were located mainly in the ventral region of the section +16.46 mm, and IN24 was in the lateral region of the section +13.45 mm (Figure 5F, H). These preferential distributions of cell types were further supported by the nucleus-specific expression of marker genes for each cell type (Figure 5G, I). The complexity in the subdomain organization and cell type composition found here demonstrates that spatial transcriptome analysis at single-cell resolution is essential for understanding the hypothalamus at the cellular and molecular level.

### Single-cell spatial transcriptome map of monkey claustrum

The claustrum is a thin, sheet-like structure located between the insular cortex and the putamen, with the distinction of having reciprocal connections with nearly all neocortical areas (Edelstein and Denaro, 2004; Jackson et al., 2018; Torgerson et al., 2015). It is implicated in a variety of brain functions, including sensory integration, attention, impulsivity, sleep, and consciousness (Goll et al., 2015; Jackson et al., 2020; Smith et al., 2020). The cellular and molecular basis for the diverse functions of claustrum remains largely unknown (Crick and Koch, 2005). Taking advantage of our Stereo-seq dataset, we have examined in detail the heterogeneity of gene expression among claustrum neurons and the spatial distribution of various neuron subtypes.

The macaque claustrum can be divided into anterior, medial and posterior regions according to the location, size and density of cells. We performed Stereo-seq analysis and 3D reconstruction for 43 coronal sections (500-μm spacing) that contained the claustrum in the monkey m1 brain (Figures 6A, S10A and Video S3). From snRNA-seq data on tissues that contained claustrum and the adjacent cortex, we identified two distinct groups of excitatory neuron clusters based on the expression of previously identified claustrum marker genes(Erwin et al., 2021) (*GNB4*, *NTNG2* and *NR4A2*; Figure S10B, C). Our snRNA-seq analysis revealed no inhibitory claustrum neurons, in line with previous reports (Erwin et al., 2021; Graf et al., 2020; Kim et al., 2016). However, our Stereo-seq data revealed a small population of claustrum neurons expressing *GAD1* or *GAD2* (Figure S10D), indicating the existence of inhibitory neurons that were not detected by the snRNA-seq method.

**Figure 6.**
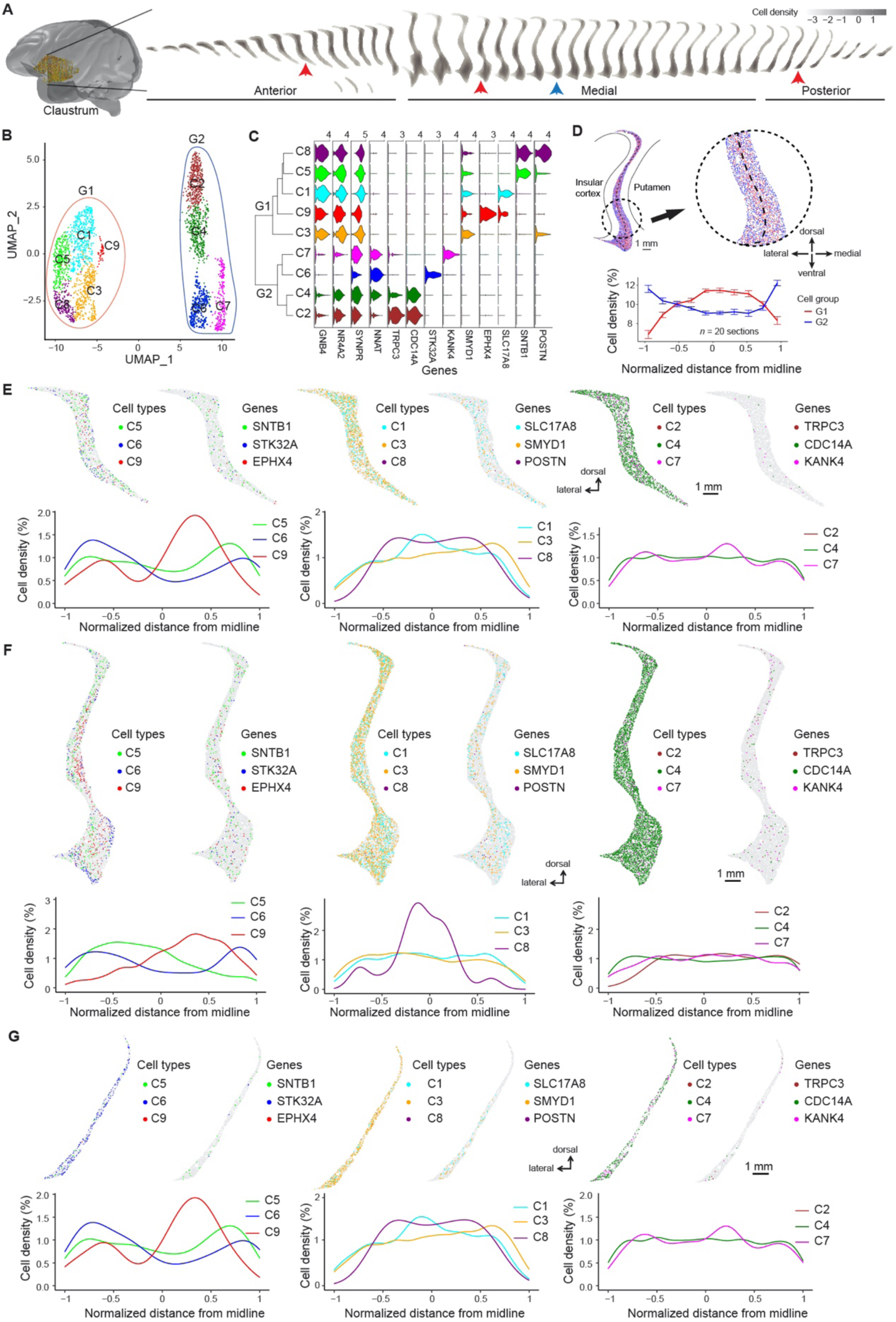
Single-cell spatial transcriptome in monkey claustrum. **(A)** Schematic diagram indicating the location of the 43 macaque brain sections containing the claustrum. The 3D movie of the claustrum reconstructed from the Stereo-seq sections in video S2. **(B)** Unsupervised clustering of snRNA-seq data from claustrum cells revealed nine cell types that fell into two groups (G1 and G2). **(C)** Log-normalized expression levels of macaque claustrum marker genes, four genes (*GNB4*, *NR4A2*, *SYNPR*, *NNAT*) were also found in mice. **(D)** Spatial distribution of G1 and G2 cells within an example medial section (red arrowhead in A). Circled region is enlarged on the right showing preferential core and shell distribution of G1 and G2 cells (marked by red and green dots), respectively. Curves below show the distribution of G1 and G2 cells, quantified by the averaged cell density (mean ± s.e.m.) at 10 bins along the normalized axis perpendicular to the midline through the section (black dashed line) for the 20 medial sections. **(E)** The spatial distribution of 9 cell types and expression of 9 example marker genes at single-cell resolution in an example section (marked by blue arrowhead in A) of anterior claustrum. Each panel showing 3 cell types and their corresponding marker genes (above), and the cell density (below) along the normalized axis perpendicular to the midline, similar to that in D. **(F-G)** As in E for example sections (marked by blue arrowheads in A) of medial and posterior claustrum, respectively.

Further clustering analysis of the claustrum cells revealed 9 excitatory neuron clusters that fell into two distinct groups (G1 and G2; Figure 6B, C). Spatial mapping of these cell types in the medial sections revealed that G1 and G2 cells were preferentially located in the core and shell regions, respectively (Figure 6D). Our results showed that the gene expression patterns in monkey claustrum showed both similarity and difference compared to those observed in mice (Erwin et al., 2021). In particular, G1 and G2 cells were marked by high-level expression of *GNB4* and *NNAT*, respectively, and were preferentially located in the core- and shell-like regions (Figure 6C), consistent with that found in mice (Erwin et al., 2021). However, *SYNPR* that uniquely marked cells in the core-like region of mice were found to be expressed in all monkey claustrum cells, and *SMYD1* gene was identified to be a new unique marker for the core-like region in the monkey. In addition, we analyzed the spatial distribution of all 9 cell types and marker gene expression patterns in all 43 sections (Figure S10E). Three representative sections at anterior, medial and posterior locations were shown at a higher resolution in Figure 6E-G, and density plots revealed both heterogeneity and homogeneity in cell distribution along the medial-lateral axis (Figure 6F). In particular, *SMYD1*-and *SNTB1*-expressing C5 and *EPHX4*-expressing C9 cells were preferentially localized to medial and lateral regions, respectively (Figures 6F and S10E). Most other cell types were found to be rather uniformly distributed (Figure 6E-G). In contrast to the hypothalamus (Figure 5), we found no distinct transcriptome-defined nucleus across the claustrum. Taken together, these Stereo-seq analyses provide an unprecedented spatial map of various cell types in the monkey claustrum.

### Topography and coordinated pattern of gene expression for NT/NM receptors and transporters

Neurotransmitters (NT) and neuromodulators (NM) are mediators of electrical and chemical communication in the brain(Luo, 2020). Specific receptors and transporters for NT/NM are essential for brain functions and serve as major drug targets for many brain diseases (Arnsten and Wang, 2016; Luo, 2020). Based on Stereo-seq data, we examined the whole-brain topography of gene expression profiles of known subunits and family members of NT/NM receptors and transporters, including those for glutamate, GABA, dopamine, acetylcholine, noradrenaline, serotonin, and histamine. We found striking variation of expression patterns in the cortical and subcortical structures for five typical coronal sections (Figure 7A). For examples, AMPA receptor *GRIA1*, NMDA receptor *GRIN1*, kainite receptor *GRIK2*, dopamine receptor *DRD2*, and serotonin receptor *HTR2C* were highly expressed in the hippocampus, motor cortex (F1), cerebellum, striatum, and choroid plexus, respectively. Systematic analyses led to the following observations on the topography and coordinated patterns of expression for these genes.

**Figure 7.**
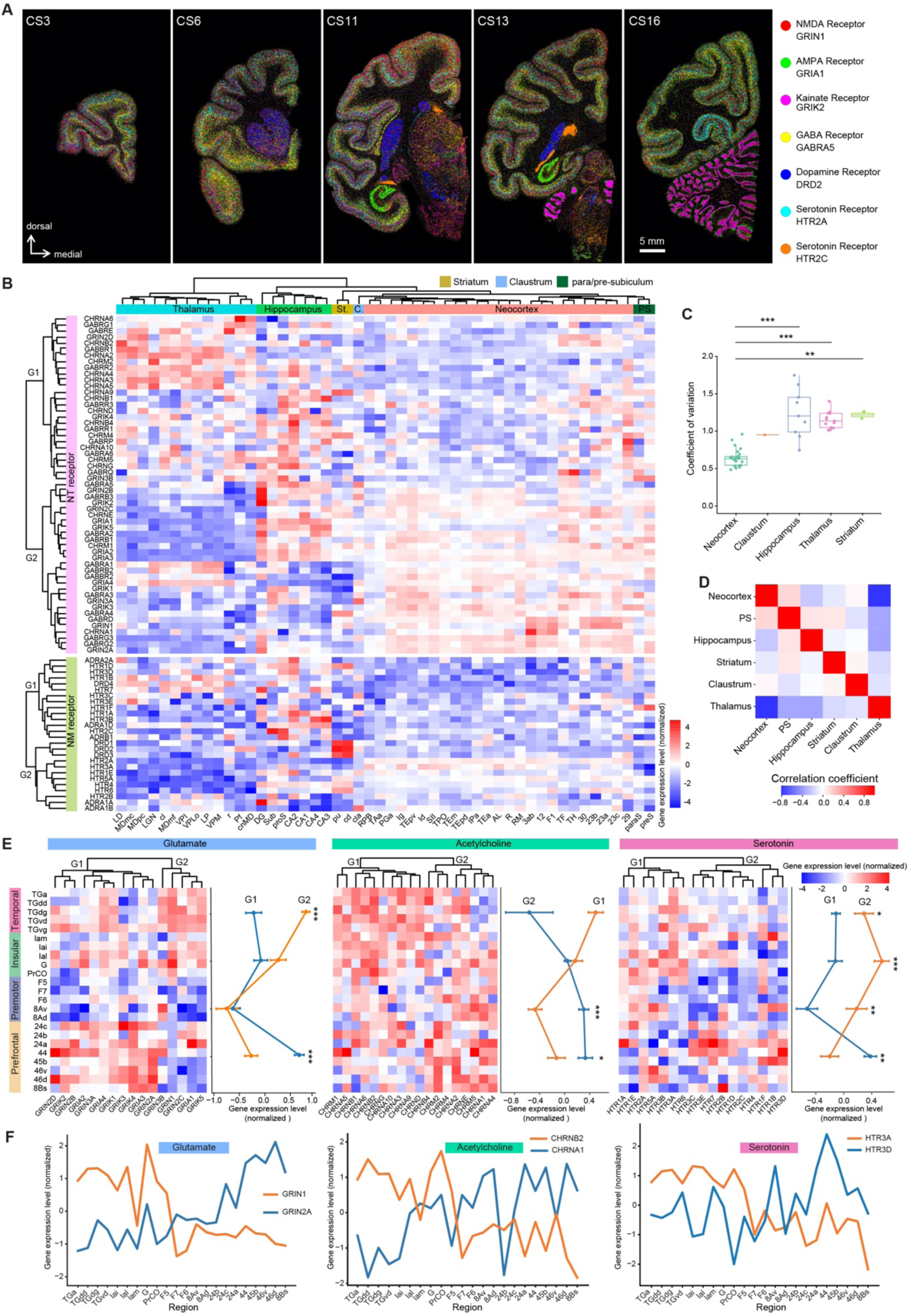
Global topography and coordinated expression of transmitter/modulator receptors. **(A)** Composite images showing gene expression profiles for 7 example NT/NM receptors in five representative coronal sections. **(B)** Heat maps of Z-score normalized expression levels of NT (top) and NM (bottom) receptor genes in coronal section CS11, with the scale shown on the right. Regions and genes were organized according to hierarchical clustering analysis of Stereo-seq data. Note that clusters of NT/NM receptor expression are distinct among areas of neocortex, thalamus, striatum and hippocampus, but similar within subregions of each of these four areas. **(C)** Variance of gene expression profiles for NT/NM receptors within subregions of the four areas in **B**. **(D)** Cross correlation of gene expression profiles for NT/NM receptors and transporters among cluster groups of the four areas in **B**. **(E)** Heat maps of Z-score normalized expression levels of NT and NM receptors for different regions of neocortex in coronal section CS6, with the scale shown on the top right. Regions were ordered by their corresponding areas. Note the opposing expression profiles between different cluster groups (G1 vs. G2), summary for expression levels of G1 and G2 cluster groups is represented by the average expression levels for all genes within the group on the right of each heatmap. **(F)** Expression level of indicated subunits of neurotransmitter genes across the temporal, insular, premotor and prefrontal cortices. Note the opposite expression profiles between NMDA receptor subunits *GRIN2A* and *GRIN1*, Acetylcholine receptor subunits *CHRNA1* and *CHRNB2*, and serotonin receptor subunits *HTR3A* and *HTR3D*.

First, we found striking differences in the expression patterns between cortical and subcortical regions, as revealed by the example genes shown in coronal sections containing both regions (Figure 7A). Hierarchical clustering analysis showed that expression patterns of these genes could be grouped into distinct clusters (CS11 in Figures 7B and S11B, other coronal sections in Figure S11A), with clusters in subcortical regions exhibiting significantly higher variation in gene expression than cortical clusters (Figure 7C). The gene expression patterns of the hippocampus were separated into two subclusters (Figure 7B and Figure S11A, B): para/pre-subiculum (referred as PS), which is spatially closer to the neocortex, showed gene expression patterns similar to those of the neocortex, whereas all other hippocampal domains were distinct from the neocortex. Thalamus and neocortex exhibited opposite expression patterns for a large group of genes for NT/NM receptors and transporters (G1 vs G2; Figure 7B). Striatum (in CS11) and cerebellum (in CS16) showed the most striking differences in gene expression from all other brain structures (Figure S11A). These observations were further supported by the strong positive correlation in gene expression profiles between PS and neocortex, and significant negative correlation between neocortex and hippocampus, striatum or thalamus (Figure 7D). Interestingly, the expression patterns in the claustrum showed no correlation with other brain areas (Figure 7D).

Second, gene expression patterns for NT/NM receptors and transporters showed positive and negative correlations within and among neocortical areas (e.g., prefrontal, temporal, insular, or premotor area), respectively (Figures 7B and S11A, C). This is in line with the notion that different regions within a neocortical area share similar circuit elements and connections. We further found that some NT/NM receptor genes exhibited high levels of expression in specific cortical layers. For examples, serotonin receptor genes *HTR2A* and *HTR2C* were highly expressed in L4 and L5, respectively (Figure S12A). Such laminar specificity of NT/NM receptor expression suggests differential synaptic composition in neurons of different layers involved in feedforward and feedback circuits (Markov et al., 2013).

Third, we found clear spatial topography of gene expression, as manifested by the expression gradient for individual NT/NM receptor subunits (Figures 7E and S11D). For examples, glutamate receptors were divided into two clusters with distinct expression patterns (G1 and G2), with G1 group showing elevated expression in the prefrontal cortex, whereas genes in G2 group were highly expressed in the temporal and insular cortices (Figure 7E, F). Specifically, genes for NMDA receptor subunits *GRIN2A* and *GRIN1* showed opposite expression gradients from the temporal, insular, premotor to prefrontal cortex (Figure 7F). Similarly, GABA receptor genes *GABRB2* and *GABRG3* showed elevated gene expression level in the prefrontal cortex, while *GABRA5* and *GABRB1* were highly expressed in a coordinated manner in the temporal and insular cortices (Figure S11D, E). The whole-brain expression patterns for NMDA receptor subunit *GRIN1* and GABA receptor *GABRA5* are shown in Figure S12B, C. Moreover, overall gene expression for acetylcholine and serotonin receptors showed opposite gradients for two coordinated groups (G1 and G2) from temporal and insular cortices to the prefrontal cortex (Figure 7E, F). The expression level of receptors for DA and NE within the prefrontal cortex was higher than that in the temporal cortex (Figure S11D).

Taken together, these analyses of NT/NM receptors and transporters exemplify the use of Stereo-seq in global mapping of the expression patterns for genes that serve specific neuronal functions in the primate brain.

## Discussion

Brain is unique among all tissues in its diversity of cell types that could be defined by distinct molecular, physiological, or morphological features. Spatial transcriptome at the single-cell resolution is an efficient approach to obtain information on both molecular fingerprints and cellular locations that are critical for determining the connectivity and functions of diverse cell types. In this study, we performed comprehensive single-cell spatial transcriptome analyses at the level of the whole monkey brain. This was enabled by the recently developed Stereo-seq technology (Chen et al., 2021) with single-cell mRNA capturing capability using nanoball-array chips with ultra-large sizes (up to 15 cm^2^). Our major findings include (1) distinct spatially resolved cell-type compositions across cortical and subcortical regions at the whole-brain level, (2) distinct cortical layer- and region-specific gene expression profiles, together with the identification of novel layer- and region-specific genes, (3) refined subregion parcellation of brain regions using gene expression patterns, (4) transcriptome-based analysis of domains and cell type compositions in several subcortical structures, hypothalamus and claustrum in particular, and (5) whole-brain topography and coordinated patterns of gene expression for NT/NM receptors and transporters. These findings provide an important foundation for understanding the spatial organization of diverse cell types and regulation of their gene expression across the whole monkey brain.

Our brain-wide transcriptome analysis was based on 20 coronal sections at different coronal coordinates across the left hemisphere of the male macaque brain, covering almost all of previously defined cortical regions (140 out of 143, as in (Saleem and Logothetis, 2007)). These data provided an initial framework for 3D spatial transcriptome atlas at the single-cell resolution of the entire macaque brain (Figure 2), which could be further refined with additional data on coronal and sagittal sections across both hemispheres. Furthermore, sexual dimorphism of gene expression patterns of the macaque brain needs to be examined. Although the number of genes captured per unit area by the present method is higher than that by previously reported methods (Chen et al., 2021), the number of captured genes per cell (663 ± 334) remained lower than that of conventional snRNA-seq method (2,990 ± 1132 in the current study). This problem of low gene number per cell was circumvented by using snRNA-seq data to identify cell types and by binning the Stereo-seq data over 25-100 μm distances (about 2-10 cell diameter). Future technical improvement in mRNA capturing capability could elevate the sensitivity and precision of the Stereo-seq technology.

We have shown that spatial transcriptome profiles obtained by Stereo-seq offer a new type of information for parcellating the brain, supplementing traditional anatomical, histological and brain imaging approaches (Saleem and Logothetis, 2007; Van Essen and Glasser, 2018). Furthermore, subregion- and layer-specific genes identified in this study could be used for further functional and mesoscopic connectome studies. Subregion boundaries, as revealed by functional brain imaging, may be dynamically regulated by experience, as shown by use-dependent expansion and shrinkage of cortical maps found in animals and humans (Buonomano and Merzenich, 1998; Elbert et al., 1995). Performing Stereo-seq analysis repeatedly at different times could allow elucidation of molecular fingerprints for experience-dependent *vs.* genetically determined regional boundaries. Compared to other species, many new regions unique to primates have emerged, such as dorsolateral prefrontal cortex (Fuster, 1997; Wise, 2012). Furthermore, brain connectivity of primates greatly differed from that of mice (Gamanut et al., 2018). Across-species comparison of spatial transcriptome will provide new molecular insights on the evolution of primate brains. Brain-wide Stereo-seq analysis of monkey models of brain diseases will also enable the discovery of novel targets for therapy.

In this study, we have identified gene expression patterns in the monkey that diverged from those in the mouse brain, even in evolutionarily ancient hypothalamus and claustrum. This supports the notion that connections and functions of these subcortical structures differ between mice and monkeys. Neocortical expansion represents an important step in the primate evolution (Fuster, 1997; Wise, 2012), and divergent gene expression patterns in subcortical structures in primates may reflect coevolution that adapt to more extensive cortical control of subcortical functions. The divergent gene expression patterns we identified in monkeys could offer targets for manipulation in studying the mechanism underlying primate-specific behaviors. Our analyses revealed two distinct groups of subcortical structures in terms of their cell type distributions, one group exhibiting clear aggregation of different cell types into subdomains with sharp boundaries (*e.g.*, hypothalamus, thalamus and hippocampus), and the other group showing much more dispersed patterns of cell-type distribution (*e.g.*, striatum and claustrum). The advantages of aggregated *vs.* dispersed cell-type distribution for establishing their connectivity and serving their functions represent an interesting direction for future investigation.

To examine whole-brain topography of functionally important genes, we focused on NT/NM receptors and transporters. We found region-specific global topography and coordinated expression patterns for subsets of these genes (Figure 7). Neocortical and subcortical regions exhibited opposite expression patterns for the majority of NT/NM receptor genes, consistent with their distinct functional differences. More NT/NM receptor genes were found to express at higher levels in the prefrontal cortex than those in insular and premotor cortices (Figure 7E-F), in line with drastic changes in neurotransmission and neuromodulation associated with the more recently evolved prefrontal cortex. Brain-wide mapping of gene expression topography could further reveal general principles governing spatial organization of other functionally important molecules in the brain. In summary, our results provide the first comprehensive brain-wide spatial transcriptome of the primate brain at single-cell resolution, paving the way for future molecular understanding of brain organization during development, aging, and behavior.

## Acknowledgements

We thank for Y-y He, L-j Lu, M-y Zhang and Y-c Sheng for their technical help in the Stereo-seq experiments, and Y-m Huang, J-k Lin, S-y Li, Y-n Sun, L-y Wang, H-x Liu and P-y Li for their help in data analysis. The project was supported by Shanghai Municipal Science and Technology Major Project (Grant No. 2018SHZDZX05), International Partnership Program of Chinese Academy of Sciences (CAS, No. 153D31KYSB20170059), Strategic Priority Research Program of CAS (Grant No. XDB32010100 and XDA27010400), the Scientific Instrument Developing Project of CAS (No. YJKYYQ20190052), the Scientific Instrument Developing Project of the National Natural Science Foundation of China (No.31827803), National Natural Science Foundation of China (No. 31900466), Lingang Lab (LG202105-01-01, LG202104-02, LG202104-02-02, LG202104-02-03, LG202104-02-04, and LG202104-02-06), and Shenzhen Basic Research Project for Excellent Young Scholars (No. 2020251518).

## Author Contributions

A.C., C.L., S.L., F.L., N.Y., K.W., M.Y., S.Z., Y.X., L.C., L.H., X.S., W.L., Z.L., H.W., Y.Z., H.Z., Q.Y., Y.W., J.P., C.L., Y.A., S.X., Y.L., M.W., X.S., S.L., Z.W., X.H., Y.Y., Y.Z., Z.L., X.T., J.L., M.Z., Y.H., R.Z., M.J., Y.L., H.Z., S.S., M.C., G.H., B.W., S.L., H.L., M.C., S.W., Q.Z., Q.Z., J.F., Z.L., Q.X., Y.H. and L.L. performed Stereo-seq and snRNA-seq experiments. Y.S., Y.L., Z.L., M.L., Z.Z., J.M., Q.S., Y.Z., H.Y., T.F., Y.L., Z.L., B.C., F.H., K.M., Z.Z., X.L., W.C., C.G., S.L., Y.H., Z.H., M.L., Y.B., S.L., B.X., and C.L. analyzed the data. J.W., H.Y., W.L., X.L., M.L., J.W., Y.L., J.H., H.X., W.H. and H.Y. provided important advice for the project and comments on the manuscript. Y.S., Y.L., M.P., L.L. and C.L. wrote the manuscript. Y.L., Z.S., L.L., Z.L., X.X. and C.L. designed and supervised the study.

## Data Availability

All raw data have been deposited to CNGB Nucleotide Sequence Archive (accession code: CNP0002035, https://db.cngb.org/search/project/CNP0002035).

## Methods

### Animal care

Animal protocol was approved (ION-2019011) by the Biomedical Research Ethics Committee of CAS Center for Excellence in Brain Science and Intelligence Technology, Chinese Academy of Sciences. Animal care complied with the guideline of this committee.

### Tissue collection

Left hemispheres were collected from three male cynomolgus monkeys (*M. fascicularis*; m1, 5-year old, 4.2 kg; m2, 6-year old, 6.7 kg; m3, 6-year old, 3.1kg). The animals were deeply anesthetized with tiletamine hydrochloride, zolazepam hydrochloride (25 mg/kg, I.M) and xylazine hydrochloride (20 mg/kg, I.M). The brain was quickly perfused at room temperature with artificial cerebrospinal fluid (ACSF, 0.6 L/kg, and perfusion speed 100 mL/min through the heart) bubbled with oxygen (influx with a mixture of 95% O_2_ and 5% CO_2_), followed with prechilled bubbled ACSF (4°C, 100mL/min). Using a stereotaxic device and a micromanipulator (SMM-200, Narishige), we coronally cut through central sulcus and divided the left hemisphere into anterior and posterior blocks. The isolated brain blocks were quickly wiped dry with sterile gauze and mixed thoroughly with 4°C OCT (4583#, Sakura). Subsequently the brain blocks were transferred to a metal mold fully embedded with 4°C OCT, quickly frozen on dry ice, and then stored in -80°C refrigerator. To minimize RNA degradation, all solutions were prepared with sterilized water containing diethyl pyrocarbonate (DEPC) (B501005-0005, Sangon Biotech), and all instruments were washed with sterilized water containing DEPC and RNase Zap (AM9780, Invitrogen). The whole tissue collection process was completed within 30 min.

### Tissue cryosection and section flattening

Before cryosection, temperature of the cooling chamber was set to -20°C. The tools involved (chip, forceps, brush and blade) were placed in the cryostat chamber in advance for pre-cooling. Immediately before sectioning, anterior and posterior blocks were placed into two separate cryostats (Thermo Fisher Cryostar NX50). At each desired coronal coordinate, cryosection was performed to obtain one 10-μm section for Stereo-seq, two 10-μm sections for immunohistochemical staining, and three-five 50-μm sections for snRNA-seq. Block face images were taken for each coordinate. Between successive days of sectioning, the tissue blocks were stored at -80°C. The thin Stereo-seq sections were firstly flattened on cold metal plane (−20°C) in cryostat by soft brush and plastic tweezers. Then the section was carefully placed onto the precooled Stereo-seq chip (−20°C). To attach a tissue section progressively on the entire chip, a Stereo-seq section was placed manually on operator’s hand to gradually raise section temperature on Stereo-seq chip. This procedure enables tissue attachment without air bubbles and tissue folding.

### Tissue-section processing for Stereo-seq

The tissue section on the Stereo-seq chip (5 cm x 3 cm or 2 cm x 3 cm) was then incubated at 37°C for 5-8 min and subsequently fixed in methanol (Sigma, 34860, precooled for 30 mins at -20°C; 40 ml methanol was added in 10 cm dish for each section) and incubated at -20°C for 30 minutes. Methanol was then dried out in a hood. Tissue section on the chip was then stained with ssDNA reagent (Invitrogen, Q10212) for 5 min and subsequently washed with 0.1x SSC buffer (Ambion, AM9770; containing 0.05 U/μl RNase inhibitor). Section images were captured using Zeiss Axio Scan Z1 microscope (at EGFP wavelength, 10-ms exposure). Tissue sections were then permeated by incubating in 0.1% pepsin (Sigma, P7000) at 37°C for 15 minutes (pepsin was prewarmed at 37°C for 3 min) in 0.01M HCl buffer (pH 2) and then washed with 0.1xSSC buffer (containing 0.05 U/μl RNase inhibitor) to remove pepsin. In this step, RNAs were released from the permeated tissue and captured by Stereo-seq chip. RNAs were then reverse transcribed for 2 hours at 42°C. After reverse transcription, tissue sections were washed with 0.1x SSC buffer and digested with tissue removal buffer (10 mM Tris-HCl, 25 mM EDTA, 100 mM NaCl, 0.5% SDS) at 55°C for 30 min, and then the chips were washed twice with 0.1x SSC buffer. The cDNA-containing chips were then subjected to Exonuclease I (NEB, M0293L) treatment for 1 hour at 37°C and were washed once with 0.1x SSC buffer. The cDNAs were amplified with Hot Start DNA Polymerase (QIAGEN). The PCR reaction protocol was: first incubation at 95°C for 5 min, 15 cycles at 98°C for 20 s, 58°C for 20 s, 72°C for 3 min and a final incubation at 72°C for 5 min. The PCR products were then purified using 0.6 x VAHTSTM DNA Clean Beads and were quantified by Qubit dsDNA HS assay kit (Invitrogen, Q32854).

### Library preparation and sequencing

One-hundred ng of cDNA (20 μl) from each sample were tagmented with Tn5 transposases (Vazyme) at 55°C for 10 mins, then the reaction was stopped by adding 5 μL of 0.02% SDS. PCR reaction mix (75 μL, Library HIFI Master Mix, Library PCR primer mix) was added to each fragmented cDNA sample. Samples were then transferred to a thermal cycler for amplification using the following protocol: 1 cycle at 95°C for 5 min, 13 cycles of tri-temperature reaction (98°C 20 s, 58°C 20 s and 72°C 30 s), and 1 cycle at 72°C for 5 min. After amplification, the PCR products were purified with 0.6x and 0.2x VAHTSTM DNA Clean Beads (VAZYME, N411-03) and were used for DNB (DNA Nano Ball) generation. Finally, the DNBs were sequenced on the DNBSEQ^TM^ T10 sequencing platform (MGI, Shenzhen, China) with 50 bp read1 and 100 bp read2.

### Immunohistochemical staining and brain region parcellation

Coronal sections adjacent to Stereo-seq chips were collected for IHC staining with NeuN antibody, which recognizes the DNA-binding, neuron-specific protein NeuN. The 10-μm sections were mounted on gelatinized glass slides, and baked to dry for 5 min at 37 °C. The mounted brain tissues were fixed with 4% paraformaldehyde in 0.1M phosphate buffer (PBS) for 10 min. After three washes in PBS, the sections were pre-incubated for 15 min in 0.5% Triton X-100 in PBS, and then 1 hour in blocking solution containing 10% normal goat serum and 0.1% Triton X-100 in PBS (0.1% PBST). Then the sections were incubated overnight at 4°C in 0.1% PBST containing the monoclonal antibody NeuN (Sigma-Aldrich, 1:1500). After three washes in PBS, the sections were incubated for 30 min in 0.1% PBST containing 0.6% hydrogen peroxide to block endogenous peroxidase that might contribute to background staining. After three washes in PBS, the sections were incubated in PBS containing biotinylated secondary antibody (1:200) for 2 hours, washed three times in PBS, and transferred to PBS containing the peroxidase conjugate from the Vectastain CBC kit (Vector Laboratories, Burlingame, CA). After rinsing three times in PBS, the sections were immersed in a solution of 0.05% 3-3’diaminobenzidine-4HCl (DAB, Sigma-Aldrich, St Louis, MO) and 0.05% hydrogen peroxide. After staining, the sections were dehydrated in increasing concentrations of ethanol, cleared in xylene and coverslipped with DPX medium. Subsequently, the glass-mounted sections were scanned at 10× (0.88 μm/pixel) in a Zeiss scanner to generate images. Based on the cytoarchitectural pattern revealed by NeuN staining and atlas-based landmarks(Saleem and Logothetis, 2007), the cortex and subcortical regions were segmented into distinct brain regions, layers, and domains.

### Brain tissue collection for snRNA-seq

These sections were cut at 50-μm thickness, and 3 to 5 sections for each coronal coordinate were collected for snRNA-seq analysis. Sections were transferred into plastic wells on dry ice and stored in a -80 °C refrigerator. Each section was further segmented into distinct areas on dry ice using tissue punchers (5 - 8 mm in diameter). Tissues at the same brain regions (e.g., around the same sulcus) were combined in a prechilled pipe as one sample. The samples were immediately frozen in liquid nitrogen and then kept in dry ice or -80 °C refrigerator. Throughout the sampling manipulation, the tissues were carefully transferred to pre-cold tube without thawing.

### Single-nucleus suspension preparation

Single nucleus suspension was prepared as previously described(Bakken et al., 2018). Briefly, frozen monkey brain tissue pieces were placed in Dounce homogenizer with 2 ml pre-chilled homogenization buffer and kept the Dounce homogenizer on ice during grinding. Tissue was homogenized with 10-15 strokes of the pestle A and followed by 10-15 strokes of the pestle B, then added 2 ml homogenization buffer to the Dounce homogenizer and filtered the homogenate through 30 μm MACS SmartStrainers (Miltenyi Biotech, #130-110-915) into 15 ml conical tube and centrifuged at 500 g for 5 mins at 4°C to pellet nuclei, then the pellet was resuspended in 1.5 ml of blocking buffer and centrifuged at 500 g for 5 mins at 4°C to pellet nuclei. Nuclei were resuspended with cell resuspension buffer for subsequent snRNA-seq library preparation.

### snRNA-seq library construction and sequencing

The DNBelab C Series High-throughput Single-Cell RNA Library Preparation Kit (MGI, #940-000047-00) was utilized to construct the sequencing libraries according to the manufacturer’s protocol. In brief, single-nucleus suspensions were used for droplet generation, emulsion breakage, beads collection, reverse transcription, second-strand synthesis, cDNA amplification and droplet index product amplification to generate barcoded libraries. The sequencing libraries were quantified by Qubit™ ssDNA Assay Kit (Thermo Fisher Scientific, #Q10212) and sequenced on the ultra-high-throughput DIPSEQ T1 or DIPSEQ T10 sequencers sequencer at the China National GeneBank (Shenzhen, China).

### snRNA-seq data processing

The sequencing data were processed as previously described(Liu et al., 2019). First, bead barcodes and unique molecular identifier (UMI) sequences were extracted using parse function in PISA (https://github.com/shiquan/PISA). For cDNA libraries, 1-10bp and 11-20bp of read1 were bead barcodes, the 21-30bp was UMI sequence, the whole read2 (100bp) was used for downstream alignment analysis. For the Droplet Index libraries, 1-10bp and 11-20bp of read1 sequences were bead barcodes, the 1-10bp of read2 was UMI sequence, the 11-20bp and 21-30bp of read2 were droplet index barcodes. Reads with improper barcodes according to the barcode list were excluded. To map the pre-mRNA fragments which may cover both exonic and intronic regions, we created a modified GTF annotation file from ensemble release-91 which only contain transcript regions, and the original annotation rows for exons were all deleted, then we replaced the feature type name from ‘transcript’ to ‘exon’. Then the snRNA-seq data were aligned to *Macaca fascicularis* genome (5.0.91) reference using STAR (v2.5.3)(Dobin et al., 2013) with the modified GTF file described above. To estimate the actual number of beads, we used the “barcodeRanks” function of DropletUtils tool to find the threshold value of sharp transition in total UMI counts distribution. Beads with UMI counts less than the threshold were removed. We merged the beads considered to be one cell(Liu et al., 2019), and counted the gene expression of cells by PISA.

We next filtered out libraries for quality control using the following criteria: 1) Reads with proper barcodes less than 1,000,000, 2) overall mapping ratio less than 85%, and 3) estimated number of cells less than 1,000 (except for 6 cerebellum libraries). As a result, 133 snRNA-seq libraries covering different brain areas were included for further downstream analysis (Table S2).

### Cell clustering and cell type identification

Basic processing and visualization of the snRNA-seq data were performed with the Seurat package (v.3.4) in R (v.4.0.5). Our initial dataset contained 855,608 cells. We discarded cells with the number of genes (nFeature) less than 200 or larger than 7000, and the percentage of mitochondrial genes (percent.mt) larger than 5%. After this quality control, 652,225 cells were included for the following analyses. The data were log normalized and scaled to 10,000 transcripts per cell. Genes with high variations across cells were identified with the “FindVariableGenes” function with default parameters. Next, principal component analysis (PCA) was carried out, and the top 50 principal components (PCs) were stored. Clusters were identified with the “FindClusters” function using the shared nearest neighbor modularity optimization with a clustering resolution set at 0.8. This method resulted in 98 cell clusters. Then, cell clusters were identified by louvain algorithm, a graph-based clustering approach. We used uniform manifold approximation and projection (UMAP) method to visualize the distance between cells in 2D space. Briefly, the value of k.param is adjusted by input cell numbers. The edges between two cells were measured by Jaccard distance to construct shared nearest-neighbor (SNN) graph. Then graph-based clustering was performed for 50 times and modularity was calculated until reaching the plateau. “FindAllMarkers” and “FindMarkers” functions were used for differential gene expression analysis between clusters. For each cell type, we used multiple cell type-specific genes that have been previously described in the literature(Ximerakis et al., 2019; Yuste et al., 2020; Zeisel et al., 2018) to determine cell-type identity.

### Correlation-based circling (CBC) method for single cell identification of Stereo-seq data

We developed a correlation-based circling method to register captured mRNAs to the corresponding single cells in Stereo-seq data. The CBC method included four steps: (1) We simulated an “ones” matrix with the area equal to a single cell (diameter 15 μm or 30 DNBs; referred to as cell mask), and then traversed the Stereo-seq gene expression matrix to calculate the correlation between total RNA expression in each square (15 μm x 15 μm) and cell mask using the “normxcorr2” function in MATLAB (version 2021a); (2) The distribution of correlations was calculated for each section, and mean + 3 standard deviation was used as a cutoff for filtering noise signals so that DNBs with expression level larger than the cutoff were set as 1 and others set as 0; (3) Then 4-connected DNBs were assigned as single cells; (4) We next filtered assigned cells with area less than 1/3 of the actual cell size (∼145 DNBs). For assigned cells with length larger than 30 μm or areas larger than 1500 DNBs, we further increased the threshold in step 2 to split these large assigned cells into smaller ones until their sizes complied with the criteria in step 4.

### Single cell annotation of Stereo-seq data

For single cells registered by CBC method from Stereo-seq transcriptomic data, we employed standard pre-processing steps similar to those for snRNA-seq datasets. Specifically, we first performed quality control to remove low-quality cells with the following criteria: (1) the number of genes detected in each cell lower than 100 or larger than 2000, (2) percentage of mitochondrial genes (percent.mt) larger than 5%. On average 180 thousand cells were included for each section after the initial quality control. We then performed log2 normalization for all the cells in each section, using a size factor of 10,000 molecules for each cell with “NormalizeData” in Seurat. Expression values for each gene were Z-score transformed across all cells with “ScaleData” function in Seurat, which is standard protocol prior to running PCA. Then “FindVariableFeatures” function was applied to the scaled datasets of both snRNA-seq and Stereo-seq data for identification of a subset of features exhibiting high variability across cells (*n* = 3000). This step resulted in heterogeneous genes to prioritize the downstream analysis.

To annotate single cells on Stereo-seq data with cell types classified by snRNA-seq analysis, we next identified anchors using the “FindTransferAnchors” function for each of the two datasets. These anchors represent two cells (with one cell from each dataset) that were predicted to originate from a common biological source. We then created a binary classification matrix containing the classification information for each anchor cell in the reference (snRNA-seq) dataset. Each row and column of the matrix corresponds to a possible class and a reference anchor, respectively. If the reference cell in the anchor belongs to the corresponding class, the entry in the matrix was filled with ‘1’. Otherwise, the entry was assigned as ‘0’. We then computed label predictions by multiplying the anchor classification matrix with the transpose of the weight matrix using the “TransferData” function of Seurat.

### Identification of layer-specific genes

We performed differential expression analysis among layers for each cortical region. First, the expression matrix of one specific cortical brain region was extracted. Next, for each layer of this region, UMI from 2500 neighboring DNBs (50 DNBs * 50 DNBs) were summed together to create bin50 expression matrix. Then, the bin50 expression matrix for each cortical layer was preprocessed using Seurat (3.2.2) “CreateSeuratObject” function and merged together using “merge” function given a layer label to create a labeled Seurat object. After that, we normalized the total UMI of every bin50 multiplied by a scale factor of 10,000 and transformed the normalized data into log space using the “NormalizeData” function. Finally, the differentially expressed genes of each layer were calculated using “FindAllMarkers” function. We used the following criteria to defined layer-specific genes: (1) Fold change ≥ 1.25, (2) adjusted *P* value ≤ 0.01, Wilcox rank sum test, (3) the percentage of bin50 spots expressing this gene in a given layer (Pct.1) more than 1%; (4) Pct.1 > Pct.2 (the percentage of bin50 spots expressing this gene in all the other layers); (5) (Pct.1-Pct.2)/Pct.1 ≥ 0.4. In total, we identified 2,638 layer-specific genes in 103 cortical areas for five representative coronal sections (Table S4).

### Identification of region-specific genes for cortical regions

We applied hidden Markov models to identify marker genes for different cortical regions. The Stereo-seq map of the neocortex in each coronal section was first flattened to cortical sheet and then divided vertically into 15-μm segments. For each gene, we used a Gaussian filter for smoothing UMI counts across segments. Then log counts per million (CPM) was calculated for each gene for each segment. A Gaussian hidden Markov model with three states were fitted for each gene, which represented high, low and fluctuating states. The state of small regions spanning less than 240 segments (∼3mm) were initially assigned as fluctuating state. Then fluctuating regions spanning less than 96 segments between two identically assigned regions were reassigned into the same high or low state of its neighboring region. An example final state assigned by the model was shown in Figure S6A.

Marker genes were then filtered using the model we trained above. First, 20% genes with the lowest expression state were discarded. Then 20% genes with the lowest fold change between high and low expression states were also removed. Then for each cortical region, we only kept genes with high expression state occupying at least 75% segments of the region. Finally, all marker genes were sorted by the differences between the mean expression level in a given cortical region and that of all other regions. Genes with fold changes > 1.5 of the difference in the coronal section were considered as significant marker genes (Table S5).

### Cortical boundary parcellation based on Stereo-seq data

The flattened cortical sheets were divided into vertical sliding segments (150-μm width, 15-μm step size), and each segment was further divided horizontally and evenly into 20 laminar sub-divisions. Expression profiles of all genes were re-gridded into profiles in the 2D flattened cortical sheet, with a 20-division vector representing gene expression for each segment. Expression gradient on the cortical sheet was then defined by calculating vectorial differences across segments for all genes. To smooth local signal variations, a sliding window of 50 segments were applied before identifying distance peaks. An overall gradient along the cortical sheet was generated by summing gradients from top 10 genes of vectorial differences. Putative gradient peaks were identified by thresholding at 2 standard deviations, and these boundaries were further visually inspected and compared with anatomically defined boundaries. Peak-defined boundaries within 500 μm of anatomical boundaries were defined as Hit. The percentage of “Hit” (“Hit rate”) was defined by the number of “Hit” boundaries divided by the total number of anatomically defined cortical boundaries. To examine the statistical significance of the Stereo-seq defined cortical boundaries, the gradient peaks were randomly permuted to generate null distribution of “Hit rate”.

**Figure S1.**
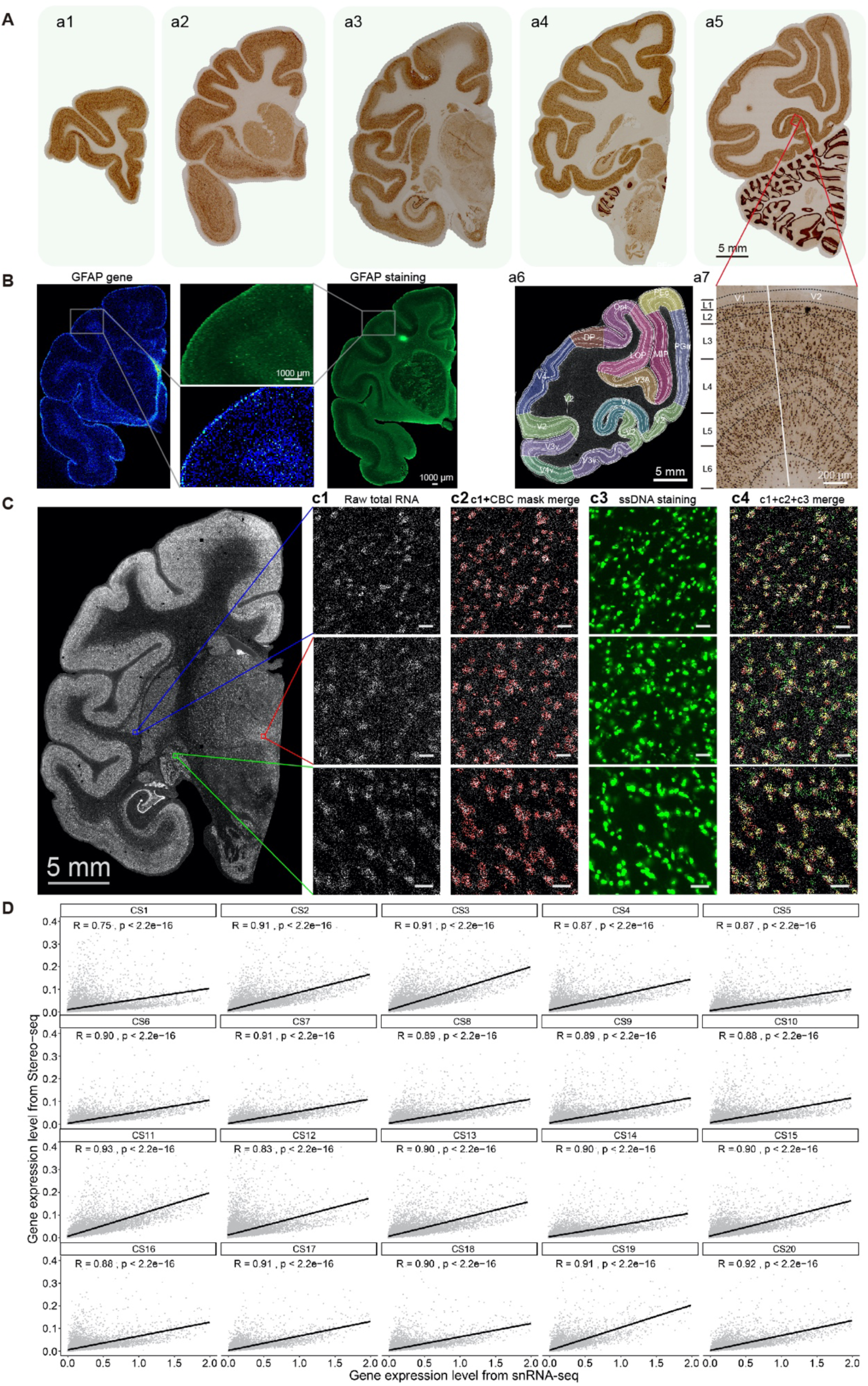
IHC staining and single-cell registration for Stereo-seq data. **(A)** Anatomical parcellation of brain regions based on IHC staining. **a_1-5_**, Images of IHC staining (NeuN) for the five coronal coordinates. **a_6_**, Example parcellation between V1 and V2 region (enlarged from the red box in a_5_), boundary (white bar) was based on IHC staining. **a_7_**, Parcellation results of CS16, colors indicate for different layers (L1-6). **(B)** Histological validation of Stereo-seq profile (left) of *GFAP* expression by IHC staining (right). **(C)** Total RNA captured by Stereo-seq of CS11. **(c_1_)** Total RNA captured for sampled locations marked by red, blue and green boxes in **c_1_**. **(c_2_)** Cells identified by CBC method for sampled locations in **c_1_**. **(c_3_)** Staining for ssDNA for sampled locations in **c_1_**. **(c_4_)** Merged image of the **c_1,_ c_2_** and **c_3_**, showing CBC based masking overlapped well with single cells stained by ssDNA. Scale bar, 20 μm. **(D)** Correlation of gene expression levels for Stereo-seq and snRNA-seq data of 20 coronal sections. Each dot represents one gene.

**Figure S2.**
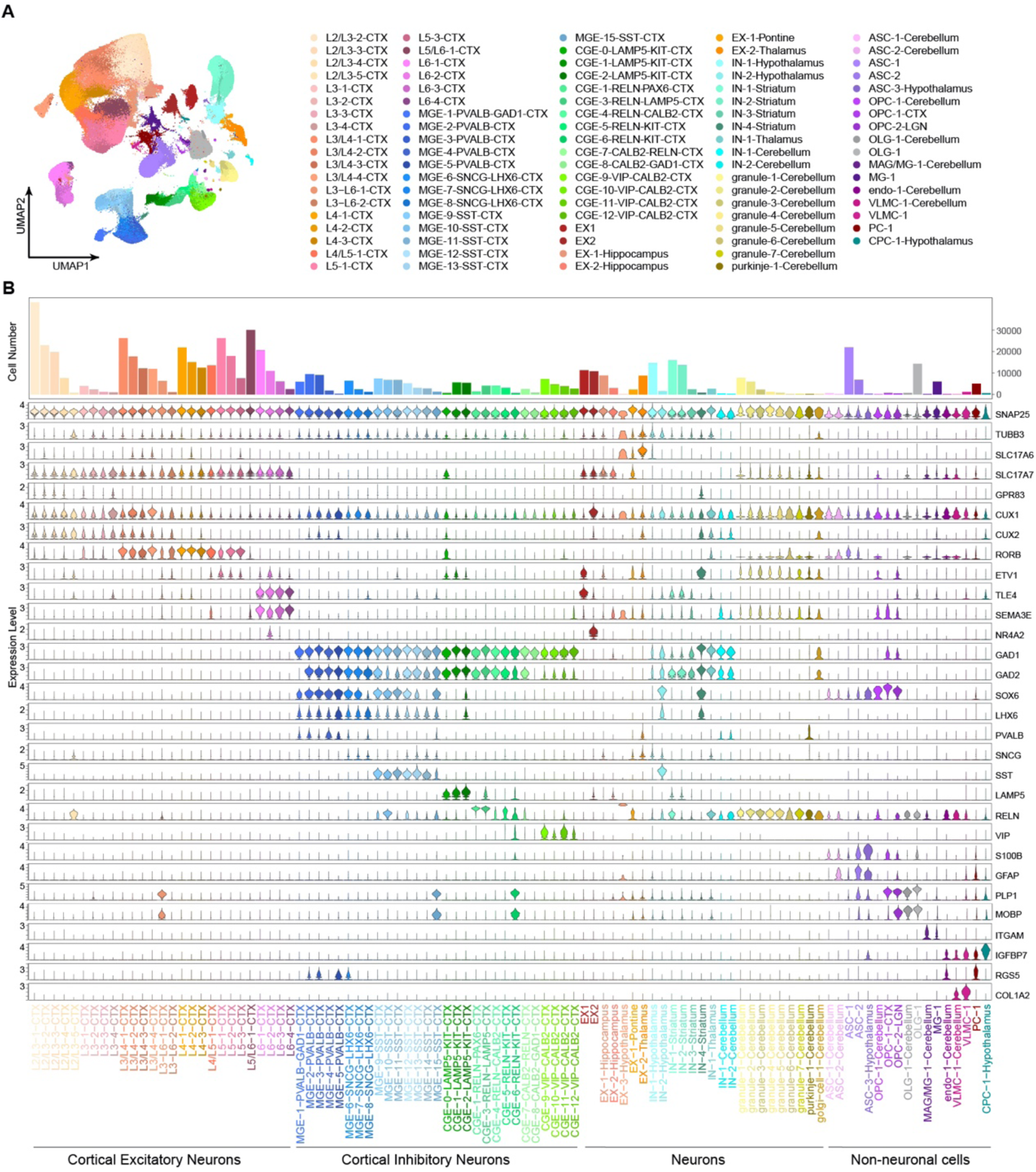
Clustering analysis and cell type annotation based on snRNA-seq data. **(A)** Unsupervised clustering analysis of snRNA-seq data on dissected tissues. Ninety-eight different cell types are coded by colors. List of cell type abbreviations see Table S3. **(B)** Example marker genes used for the annotation of cell clusters from snRNA-seq data. The top panel showed the total cell numbers of each cell type. The 98 cell types were grouped into 4 big classes based on the expression of neuronal markers and region sources. Complete list of marker genes of each cell type is shown in Table S3.

**Figure S3.**
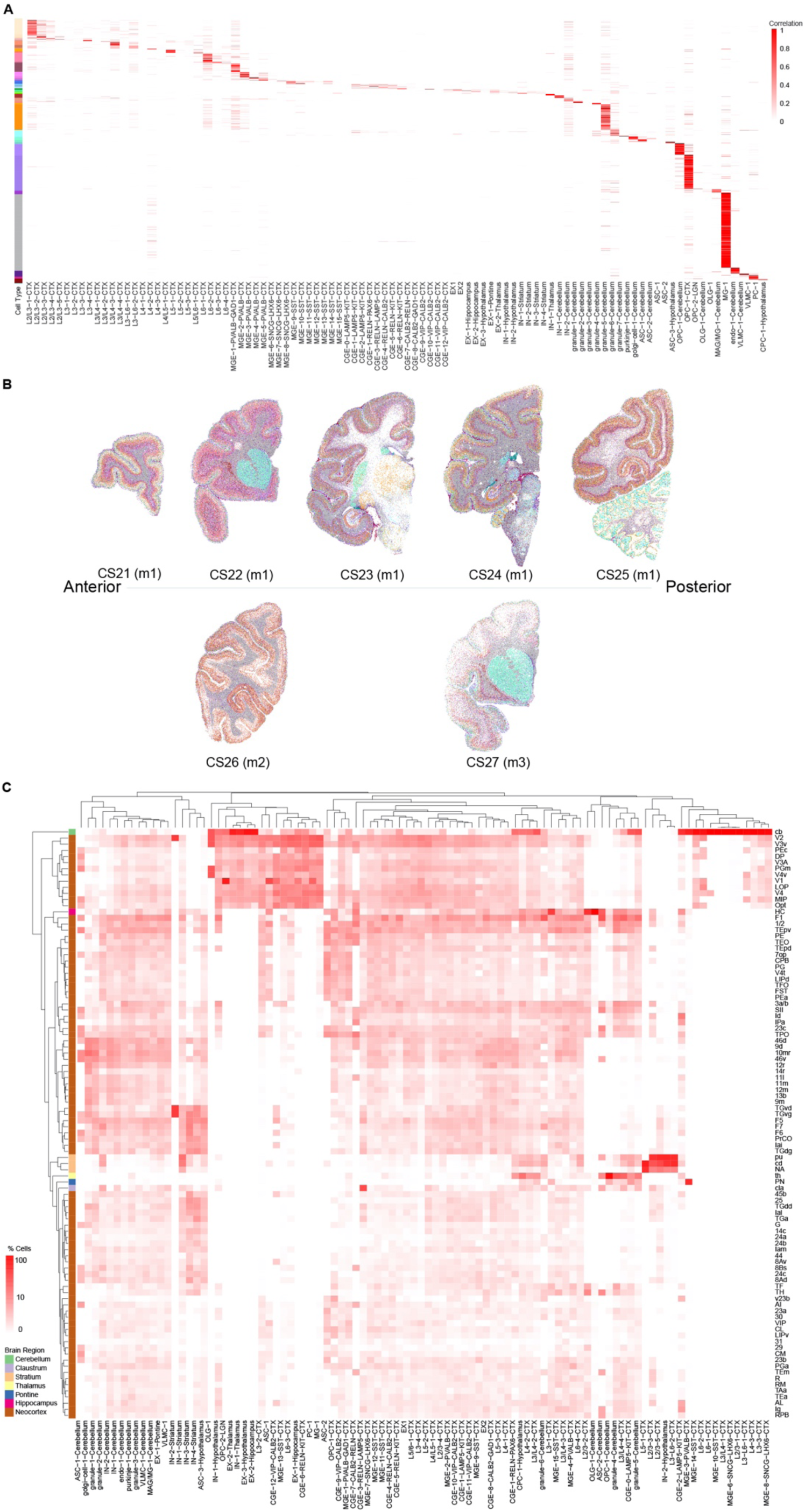
Cell type annotation of Stereo-seq data based on snRNA-seq clustering results. **(A)** The magnitude of correlation coefficient (CC) of gene expression profiles of Stereo-seq data and snRNA-seq data for each cell type (see Methods), including data for 20 coronal sections. Colors on the left coded for cell types presented as in Figure S2A. **(B)** Images depicting spatial map of snRNA-seq annotated cell types, for C21-27, as in Figure 1E. **(C)** Cell type composition over the whole monkey brain revealed by combined Stereo-seq and snRNA-seq analyses, including data for various neocortical and subcortical regions. Brain regions and cell types were ordered based on the hierarchical clustering results. The intensity of colors depicts the percentage of cell types within each brain region.

**Figure S4.**
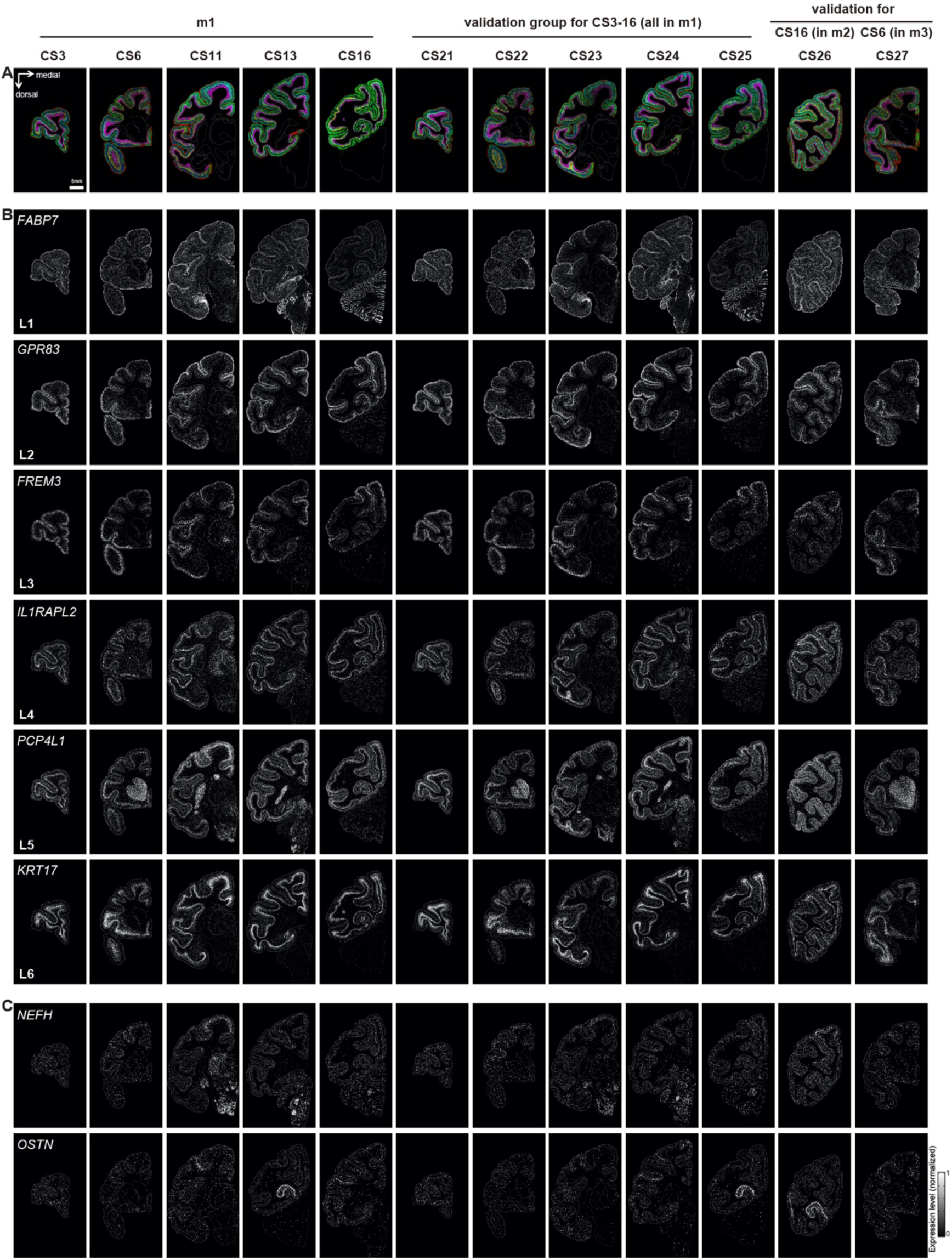
Spatial expression profiles of example layer-specific genes. **(A)** Composite image of expression profiles of 6 example layer-specific genes across the representative coronal sections from m1-3, with each gene coded with a distinct color and its expression level (summed in 25-μm bins) coded by color intensity. CS21-25 were validation sections in m1, each section at 500-μm distance from CS3, 6, 11, 13 and 16, respectively. CS26 and CS27 were validation sections for CS16 and CS6 in m2 and m3, respectively. **(B)** Six layer-specific genes that showed similar specificity in different brain regions across 12 coronal sections from m1-3. **(C)** Two layer-specific genes that showed layer specificity restricted to distinct regions across 12 coronal sections from m1-3.

**Figure S5.**
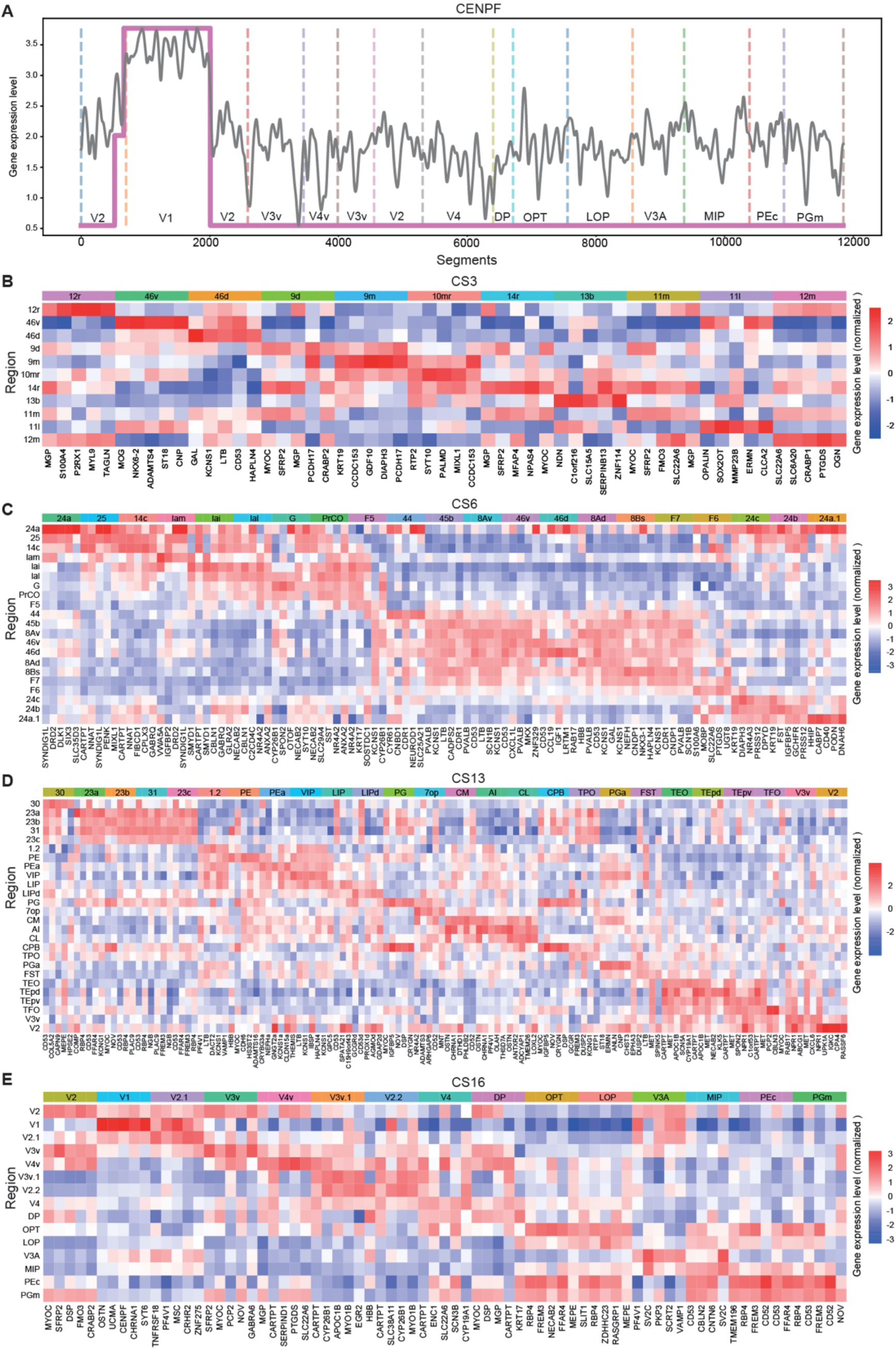
Region-specific gene expression profiles in the neocortex. **(A)** Example of log2 expression level (grey curve) for a region-specific gene *CENPF* across the vertical segments along the cortical sheet in CS11. Each segment represents 15μm-bin along the cortical sheet. Pink line marked the fitting result for Hidden Markov model (see Methods). **(B-E)**, Heatmap of region-specific gene expression levels in representative coronal sections CS3, 6, 13 and 16. Full names for region abbreviations shown in Table S1.

**Figure S6.**
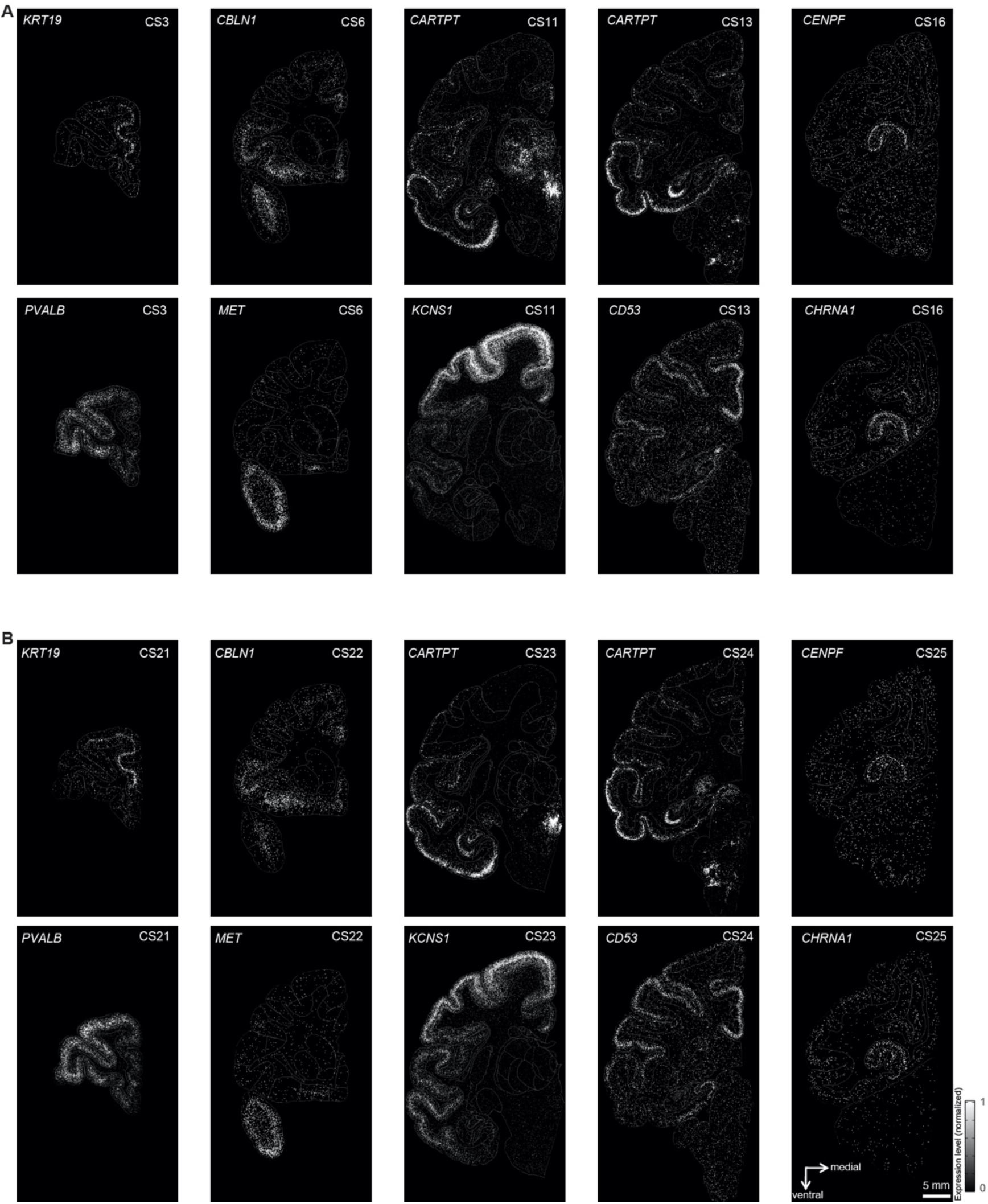
Spatial expression profiles for example region-specific genes. **(A)** Example region-specific genes for five representative coronal sections. **(B)** Validation of gene expression profiles for the adjacent coronal sections of m1. CS21-25 were validation sections, each section at 500-μm distance from CS3, 6, 11, 13 and 16, respectively.

**Figure S7.**
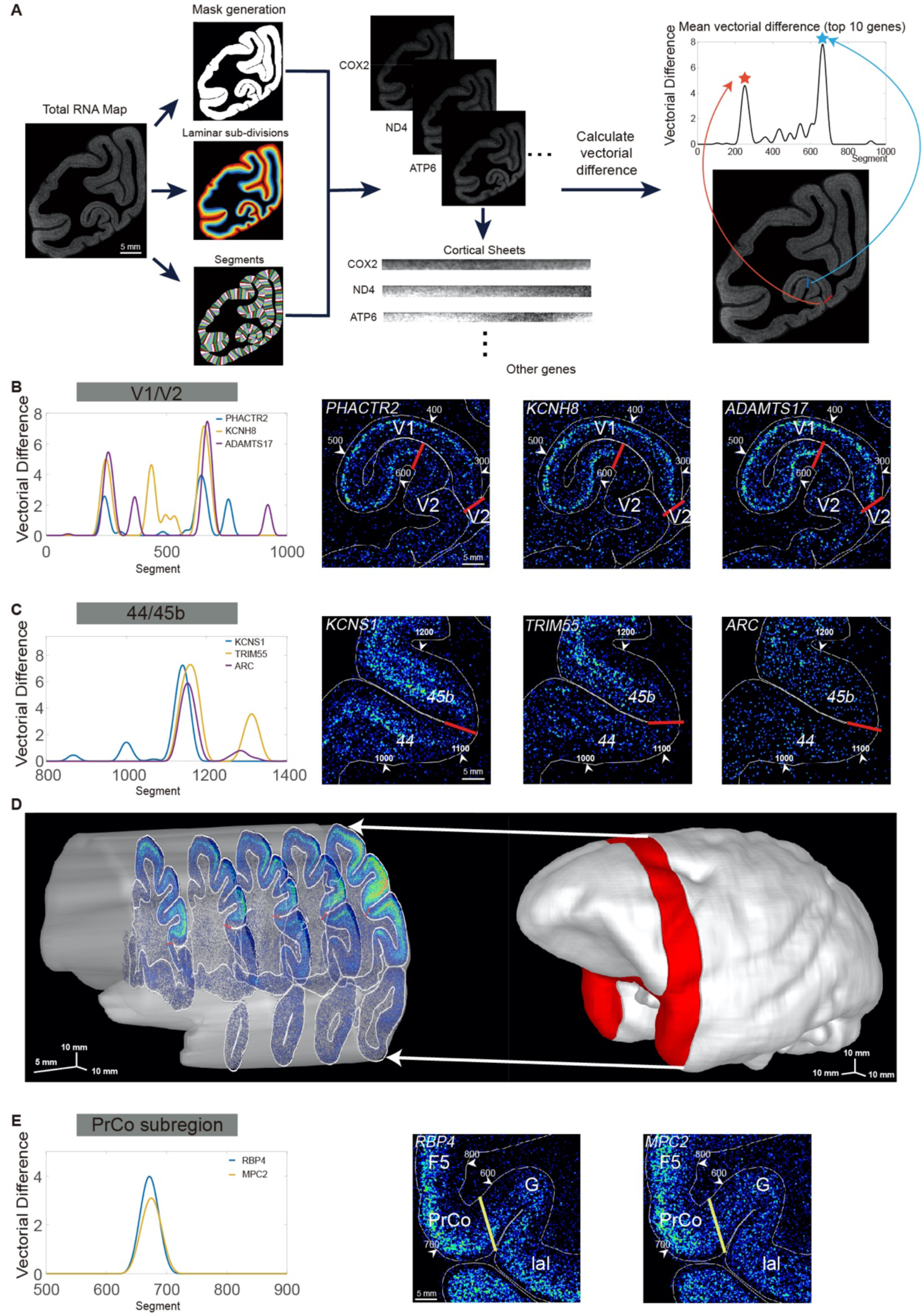
Overview of region boundary identification based on Stereo-seq data. **(A)** Schematic illustration showing the procedure for region boundary identification and parcellation (see text and Methods). The cortical sheets were divided into vertical sliding segments (150-μm width, 15-μm step size), with each segment divided horizontally and evenly into 20 laminar sub-divisions (“mask generation”). The expression level of each gene within a segmental mask was represented by a 20-dimension vector, with the component representing the average expression level of the gene within each laminar sub-division. The 20-dimension vector was used to calculate the vectorial difference between adjacent segments along the cortical sheet for each gene. Red and blue bars indicated the identified boundaries corresponding to the peaks of the vectorial difference curve summated for top 10 genes (marked by stars). **(B)** Spatial expression patterns of three additional genes mark the boundaries of V1/V2, corresponding to that described in Figure 3E-F. **(C)** Regional boundary identification between 44 and 45b using gene expression profiles. Left curves represent vectorial differences of expression profiles of 3 genes (*KCNS1, TRIM55* and *ARC*) along the cortical sheet, with their spatial expression pattern shown in the images on the right. Red lines indicate the 44/45b boundaries. **(D)** Anterior-posterior elongated 3D map showing 44/45b boundaries across 5 coronal sections (500-μm spacing) of the m1 monkey brain. **(G)** Spatial expression patterns of two additional genes marked the new boundary in the PrCO region, corresponding to that described in Figure 3H.

**Figure S8.**
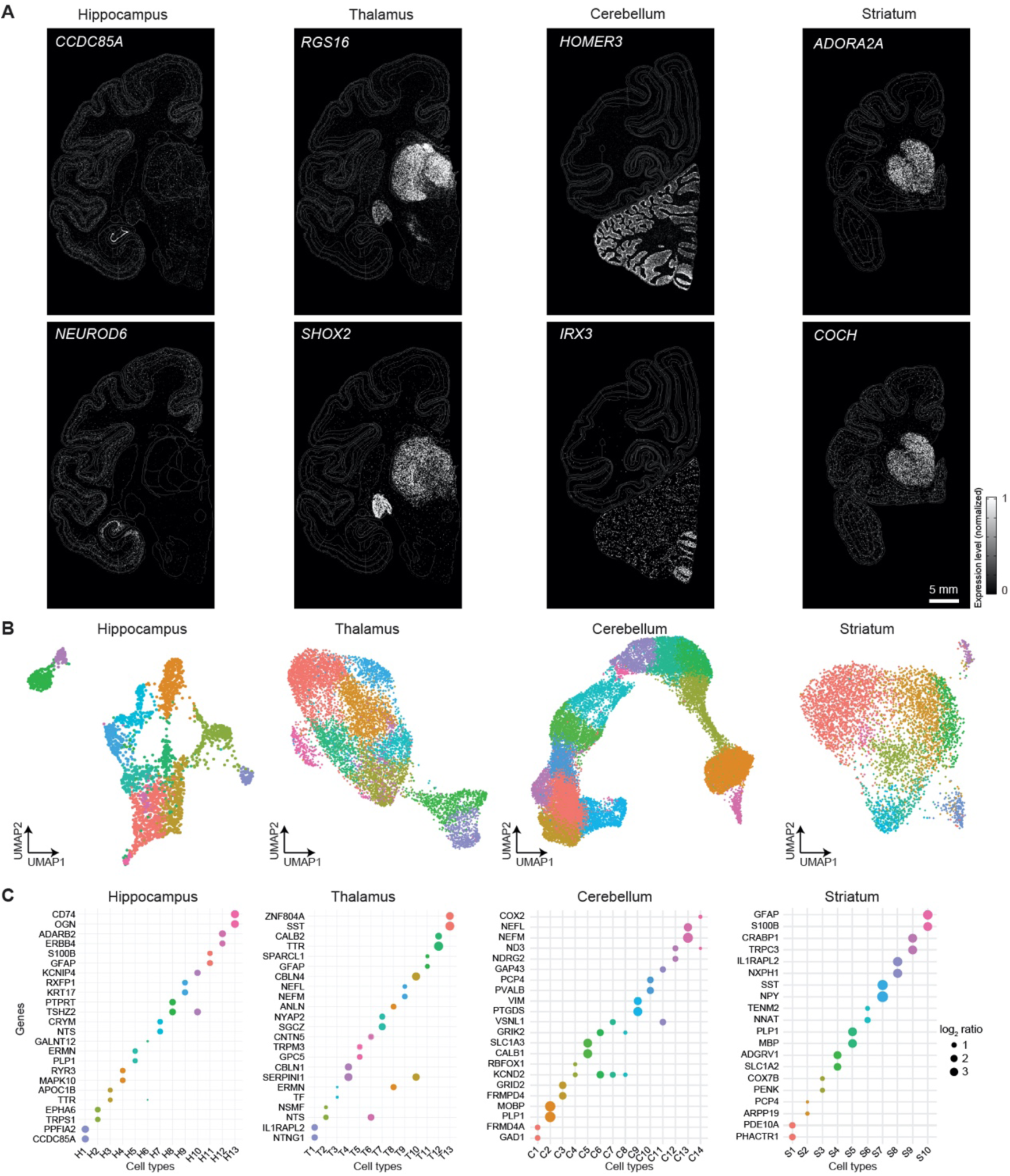
Spatial expression profiles of example genes in some subcortical structures. **(A)** Example genes that expressed at higher levels in the hippocampus, thalamus, cerebellum and striatum. **(B)** UMAP plots showing the clustering of gene expression profiles based on Stereo-seq data at 100-μm bins for the four subcortical regions. **(C)** Marker genes (two for each cluster) identified in each of the four subcortical regions. Dot size represents the log2 ratio of gene expression level in the cluster relative to the average expression level in all other clusters.

**Figure S9.**
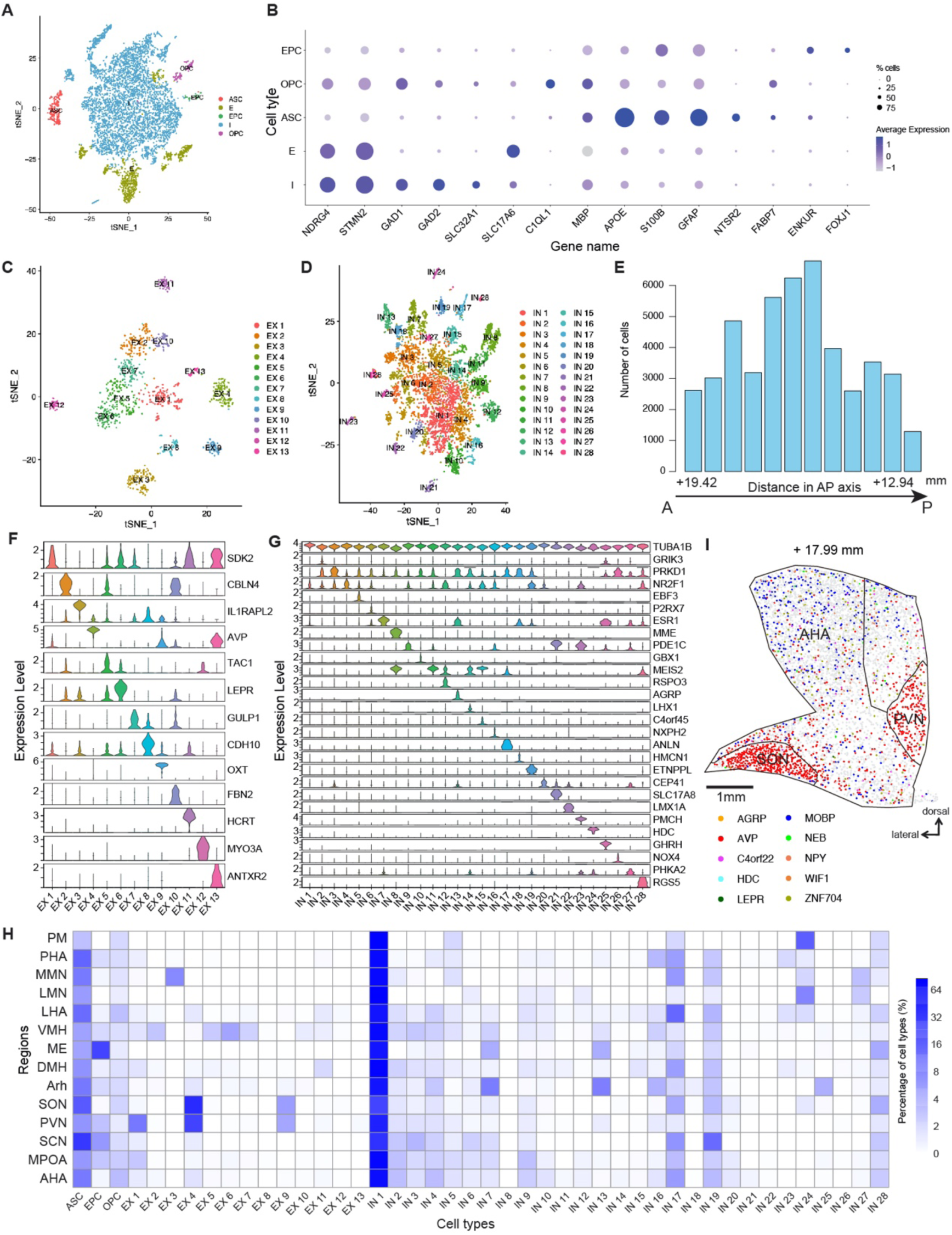
Supporting data for the spatial distribution of various cell types in the hypothalamus. **(A)** Unsupervised clustering analysis of hypothalamic snRNA-seq data. **(B)** Percent of cells expressing the specific gene (dot size) and the expression level (color intensity) for selected marker genes of indicated cell types identified by snRNA-seq data of hypothalamus tissues. **(C)** Further clustering of hypothalamic excitatory neurons into 13 subtypes. **(D)** Further clustering of hypothalamic inhibitory neurons into 28 subtypes. **(E)** The number of hypothalamic cells in each coronal section containing the hypothalamus. **(F)** Expression levels for marker genes of 13 excitatory neuron subtypes. **(G)** Expression levels for marker genes of 28 inhibitory neuron subtypes. **(H)** Heatmap showing the distribution of all identified hypothalamic cell types in various nuclei of the hypothalamus. Full names for abbreviations shown in Table S1. **(I)** The spatial map of marker gene expression (at single-cell resolution) in one example coronal section.

**Figure S10.**
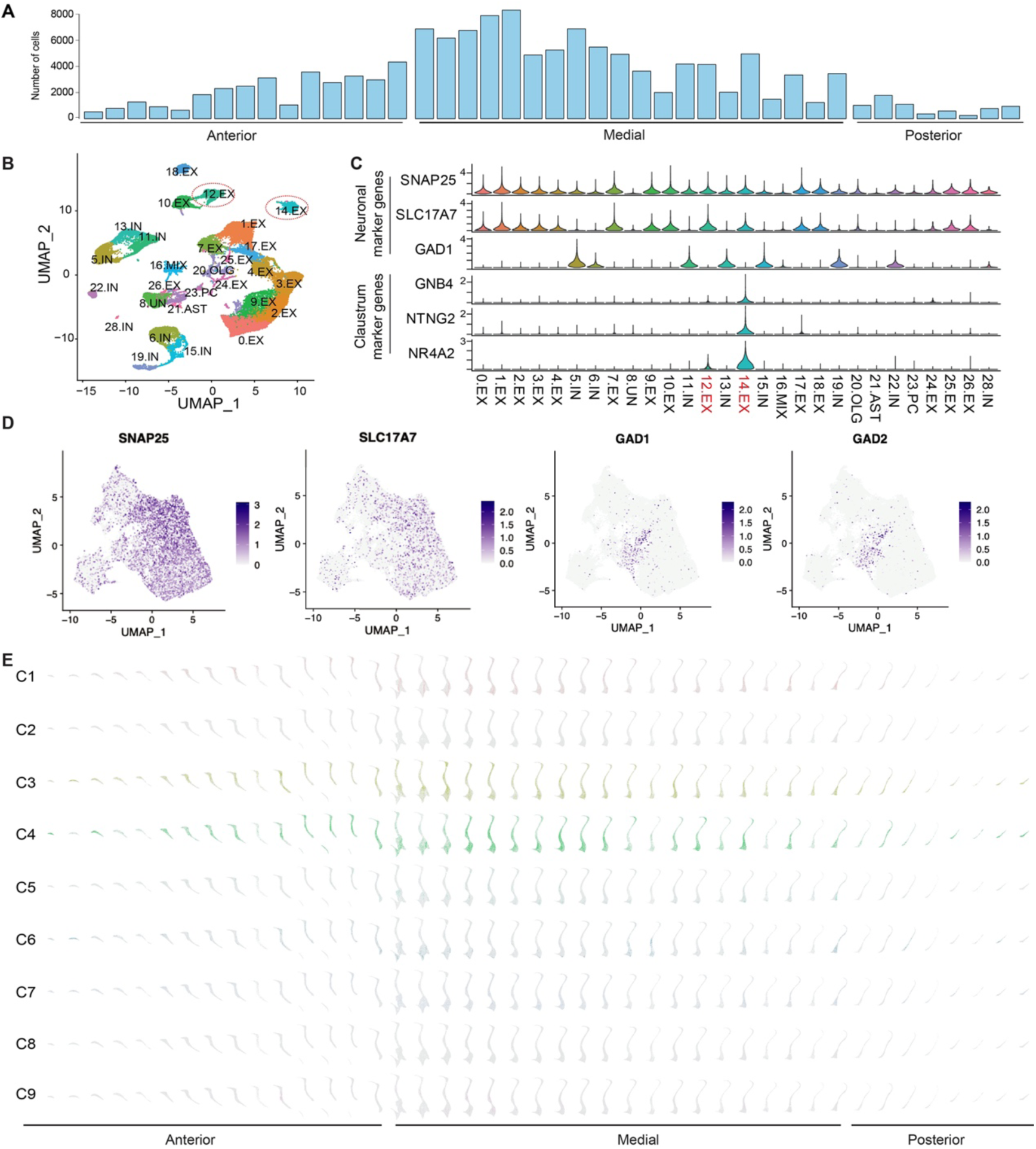
Spatial distribution of different cell types and gene expression in the monkey claustrum. **(A)** The number of claustrum cells in each coronal section containing the claustrum. **(B)** Unsupervised clustering of snRNA-seq data on dissected cortex and claustrum tissues identified 28 clusters, among which two clusters (red circled) exhibited known claustrum marker genes (see C) reported in mice. **(C)** Expression distributions of selected marker genes for the cortex and claustrum, names of claustrum-specific cell types labeled in red. **(D)** The expression of excitatory and inhibitory neuron marker genes in single cells of Stereo-seq data. **(E)** The spatial map of 9 cell types in all the claustrum coronal sections, with the grey level represents the cell density.

**Figure S11.**
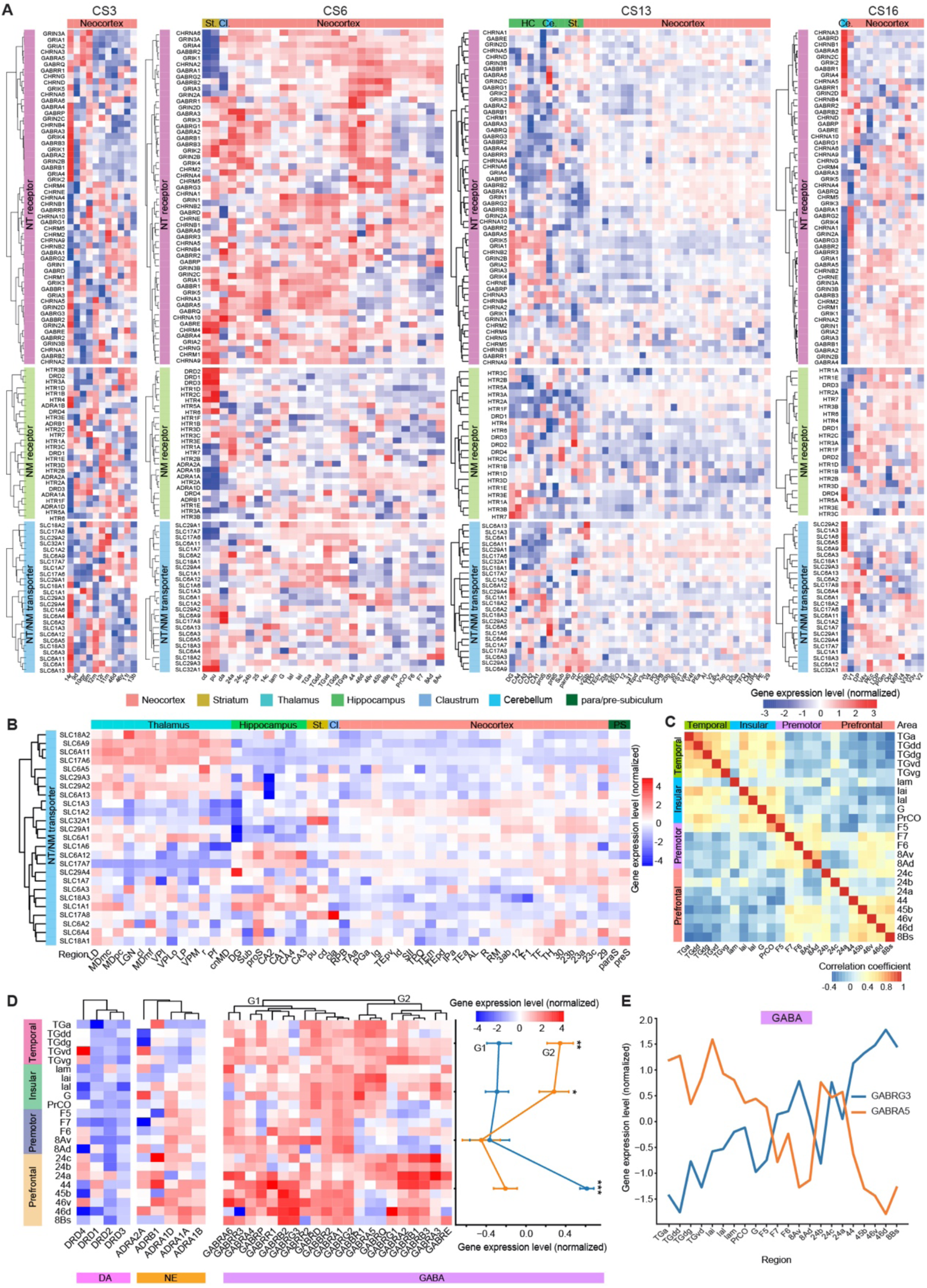
Supporting results for global topography and coordinated expression of transmitter/modulator receptors and transporters. **(A)** Heatmaps showing the expression patterns across cortical and subcortical regions for genes encoding transmitter/modulator receptor and transporters in CS3, 6, 13 and 16. Genes were organized based on hierarchical clustering results. **(B)** Heatmaps showing the expression patterns across cortical and subcortical regions for genes encoding transmitter/modulator transporters in CS11. Genes were organized based on hierarchical clustering results. **(C)** Cross correlation of gene expression profiles for NT/NM receptors and transporters among cortical regions in CS6. **(D)** Expression levels for receptors of GABA, dopamine (DA) and norepinephrine (NE) in regions of temporal, insular, premotor and prefrontal cortices for coronal section CS6. Note the opposing expression profiles between different cluster groups (G1 vs. G2) of GABA receptor units, summary for expression levels of G1 and G2 cluster groups is represented by the average expression levels for all genes within the group on the right of the heatmap. **(E)** Expression level of indicated subunits of neurotransmitter genes across the temporal, insular, premotor and prefrontal cortices. Note the opposite expression profiles between GABA receptor subunits GABRG3 and GABRA5.

**Figure S12.**
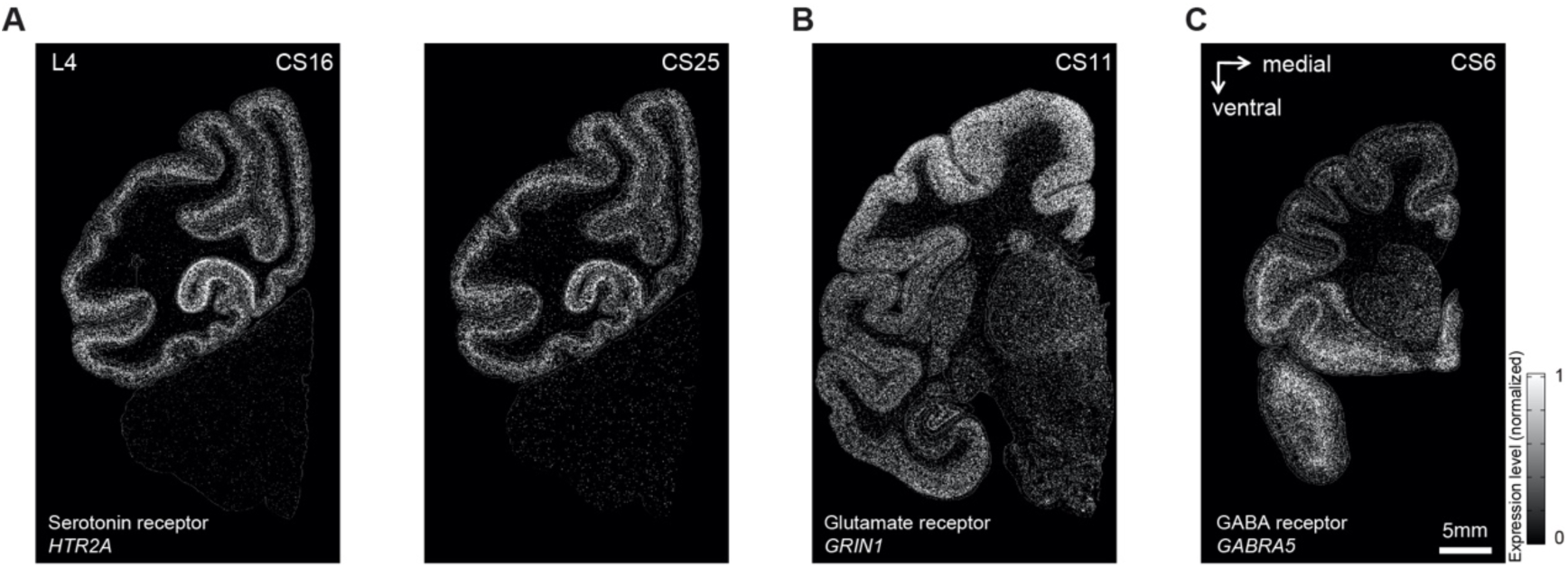
More examples showing spatial expression patterns of NT/NM receptors and transporters across the monkey brain. **(A)** Spatial expression patterns of serotonin receptor gene *HTR2A* in CS16 and CS25. **(B)** Spatial expression patterns of *GRIN1* in CS11. **(C)** Spatial expression patterns of GABA receptor *GABRA5* in CS6.

**Table S1. Full names for abbreviations of regions in this study**

**Table S2. Summary of cells and genes obtained for each library of snRNA-seq data**

**Table S3. Distribution of cell types in each cortical and subcortical structure in five coronal sections**

**Table S4. Summary of layer-specific genes in five coronal sections.**

**Table S5. Averaged expression levels of region-specific genes in each region of five coronal sections.**

**Table S6. Summary of marker genes for subdomains of subcortical regions**

**Table S7. Full names for abbreviations of neuropeptides analyzed in the hypothalamus**

